# Fast efficient coding and sensory adaptation in gain-adaptive recurrent networks

**DOI:** 10.1101/2025.07.11.664261

**Authors:** Arthur Prat-Carrabin, Maximilian V. Harl, Samuel J. Gershman

## Abstract

As the statistics of sensory environments often change, neural sensory systems must adapt to maintain useful representations. Efficient coding prescribes that neuronal tuning curves should be optimized to the prior, but whether they can adapt rapidly is unclear. Empirically, tuning curves after repeated stimulus presentations exhibit “adapter repulsion”, whose underlying mechanism remains uncertain, and which contrasts with the “prior attraction” expected under many efficient-coding models. We propose a gain-adaptive, recurrent sensory network model in which gains optimize a novel efficient-coding objective balancing accuracy and spiking cost. From the propagation of modulated gains throughout the network emerge quickly adaptive tuning curves. The model accounts for subtle adapter-repulsion effects under peaked priors and predicts fast prior attraction under broader distributions, for which we provide supporting behavioral evidence. Our framework reconciles seemingly contradictory adaptive phenomena, under a unified theoretical and mechanistic model of efficient coding mediated by gain modulation in recurrent circuits.

---

Since Barlow’s seminal proposal [1], a prominent idea in neuroscience is that sensory systems are efficiently adapted to the statistics of the stimuli that they represent [2–12]. In many models of neural sensory encoding, such ‘efficient coding’ is achieved by optimally matching neuronal sensitivities to the prior distribution of stimuli, with more encoding resources allocated to the more frequently encountered stimuli. Typical models of efficient coding posit a distribution of stimuli that is unchanging (perhaps learned over long timescales), producing static encoding schemes. Yet in natural environments the statistics of sensory inputs are not fixed: instead, they shift over time and across contexts. For an organism to maintain accurate perception, its sensory encoding must adapt accordingly, on a relatively short timescale. If the brain is able to quickly implement efficient coding — what we refer to as ‘fast efficient coding’ — then the tuning curves of sensory neurons should shift towards the most frequent stimuli, when the relative frequencies of stimuli change; a behavior we call ‘prior attraction’, and for which a recent fMRI study provides empirical support [11]. At the same time, a large body of experimental work has reported shifts in tuning curves in the seemingly opposite direction: after repeated exposure to a stimulus (the ‘adapter’), neuronal tuning curves often shift *away* from it, i.e., away from the stimulus that has become more frequent [13–18]. This behavior, which we call ‘adapter repulsion’, is moreover accompanied by several additional effects; for example, tuning curves close to the adapter typically become wider, while those further away become narrower [13–15]. Adapter repulsion occurs over durations on the order of tens of milliseconds, suggesting that neural encoding can be reconfigured fairly quickly.

These observations raise several questions. First, what mechanism underlies the fast modulation of neuronal tuning curves? Second, could the brain leverage this flexibility to implement fast efficient coding, through the prior attraction of tuning curves? Third, why then should we observe adapter repulsion (and the related effects), and how can it be reconciled with prior attraction?

Studies addressing adapter repulsion have generally attributed shifts in neuronal sensitivity to changes in synaptic weights, typically through short-term synaptic depression [13–15, 17–20]. But the implied timescales and exponentially-decaying dynamics seem ill-suited for the demands of efficient coding in dynamic environments, which needs to be both fast and sustained as long as necessary.

An alternative proposal is that adaptation is an emergent property of recurrent neural networks, even without synaptic plasticity [21, 22]. Duong *et al*. (2023) [22] show that tuning shifts can arise from the modulation of neuronal gains in a recurrent network. Inspired by this approach, we introduce a recurrent network model whose neuronal gains optimize an efficient-coding objective, formulated as the sum of a lower bound on the decoding error and a cost proportional to the neuronal spiking activity. We derive analytical approximations for the resulting tuning curves, and for the optimal gains when the prior is wide (in comparison to the width of the tuning curves). When the priors are relatively broad, the neurons’ locations (preferred stimuli) under optimal gains shift towards frequently encountered stimuli — thus exhibiting prior attraction. The same optimization framework, when applied to sharply peaked priors (corresponding to the case of a repeated adapter), yields gain profiles that deviate from the prior’s peak, with maximal gains located away from it. This distinct gain structure induces repulsion from the adapter, producing tuning shifts consistent with empirical reports of adapter repulsion, but also capturing the subtle related effects, such as the widening of tuning curves near the adapter and their narrowing further away. Our results unify prior attraction and adapter repulsion as outcomes of a single efficient coding principle, achieved solely through rapid gain modulation, without changes in synaptic connectivity.

The paper is structured as follows. First, we present behavioral evidence, obtained in a pre-registered experiment, showing that human observers rapidly adapt their sensory encoding in response to changing priors, supporting the hypothesis of fast efficient coding. We then provide schematic illustrations of prior attraction and adapter repulsion. The core of the paper focuses on the gain-adaptive recurrent-network model that we propose as a candidate neural mechanism underlying these adaptive phenomena. We show how, through the modulated gains and the recurrent interactions, a neuron’s effective location emerges as a weighted average of the locations of all the network’s neurons. We then examine decoding within this framework, and derive a lower bound on the expected decoding squared error. Next, we formulate an efficient-coding optimization problem and derive its analytical solution in the case of wide priors. This solution yields prior attraction. Turning to narrowly-peaked distributions, we show that the optimal gain configuration in this scenario places maximal gains at some distance from the peak, yielding adapter repulsion. Together, our results provide a mechanistic understanding of how sensory systems may dynamically optimize neural encoding in changing environments, and reconcile two seemingly contradictory adaptive patterns (prior attraction and adapter repulsion) as complementary outcomes of a single efficient-coding principle.

## Results

### Behavioral evidence of fast efficient coding

Before presenting in detail our model of gain-adaptive recurrent network, we exhibit some behavioral evidence of adaptation of encoding to changes in the prior. We conducted a pre-registered, online numerosity-estimation experiment. On each trial, subjects were briefly presented (for 500 ms) with a cloud of dots (Fig. 1A) and then asked to report, using a slider, their best estimate of the number of dots in the cloud. This number was chosen randomly on each trial from one of three uniform prior distributions: the Narrow prior ranged from 50 to 70 dots; the Medium prior, from 40 to 80 dots; and the Wide prior, from 30 to 90 dots. Importantly, the prior was randomly selected at every trial (with equal probabilities). Subjects were informed of the chosen prior at the onset of each trial (this information was displayed for at least one second, before subjects could make a response). This experiment is a variant of a previous one in which the trials with the same prior were presented together, in three separate blocks of trials [10]. In this original experiment, the imprecision in subjects’ responses increased with the prior width. The current experiment tests whether this adaptation also occurs on a trial-by-trial basis, when the prior is changed at every trial.

**Fig. 1:**
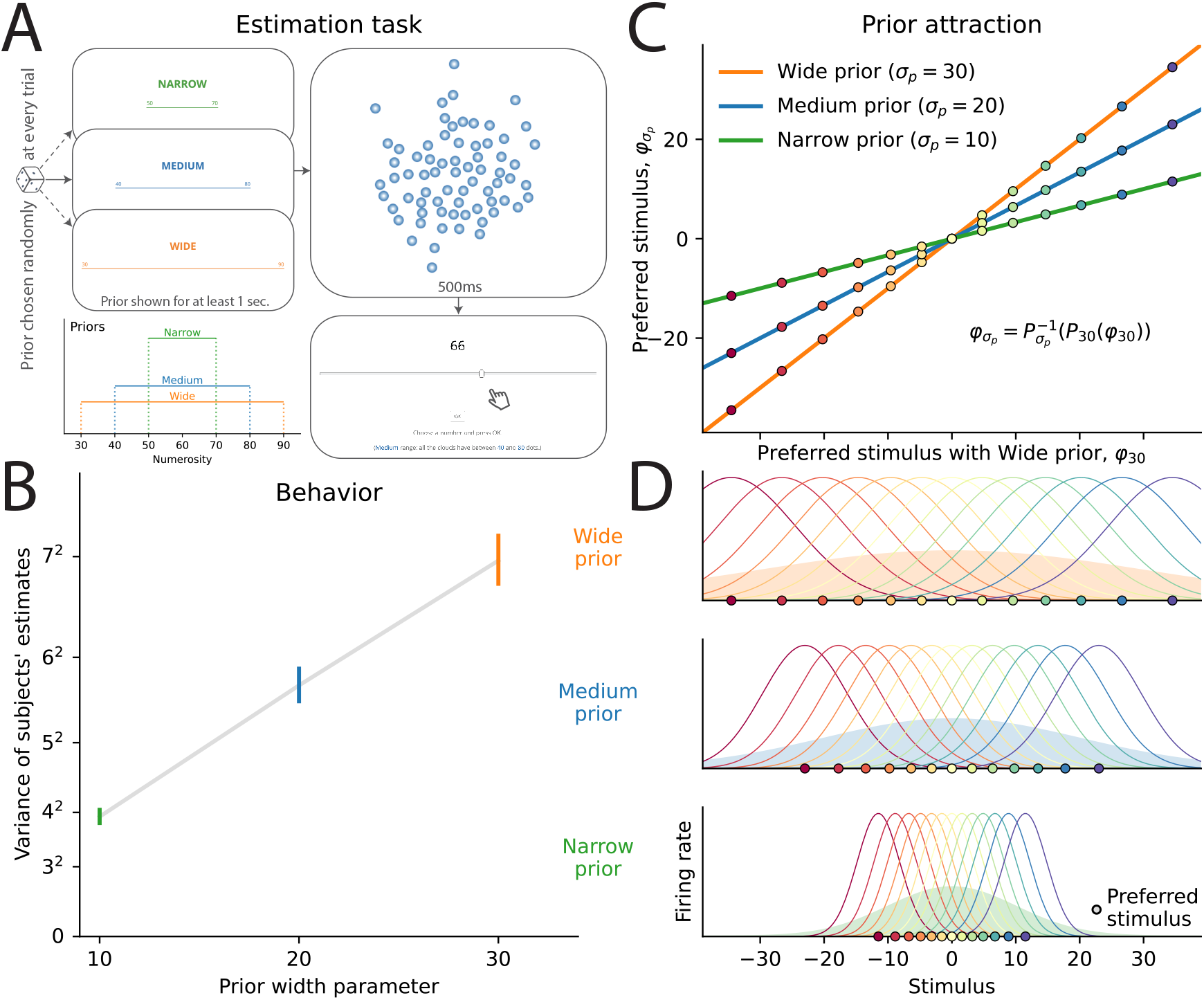
Fast efficient coding: behavioral evidence and illustration of prior attraction. **A**. Numerosity estimation task. At each trial, a uniform prior (Narrow, Medium, or Wide) is randomly chosen, and explicitly shown for at least one second and until the subject presses the spacebar. A cloud of dots is then presented for 500ms, after which the subject is asked to estimate its numerosity. **B**. Variance of subjects’ estimates as a function of the width of the prior. Values shown are Bayesian posterior mean estimates (see Methods). Error bars show the 5%-95% credible intervals. The estimate variance appears to be an affine function of the prior width. **C**. Neurons’ locations 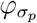 under efficient coding with each of the three priors, as a function of the locations with the Wide prior, φ_30_. The Wide line is thus the identity line; the contraction of tuning curves with narrower priors results in the gentler slopes of the other lines. Specifically, efficient coding predicts that a neuron encodes for the same quantile, under all priors, i.e., 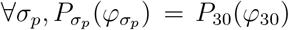, where 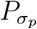 and *P*_30_ denote the priors CDFs. **D**. Prior attraction: illustration of the tuning curves predicted by an efficient-coding model, for three Gaussian priors of mean 0 and standard deviations *σ*_*p*_ equal to 10 (Narrow; bottom), 20 (Medium; middle), and 30 (Wide; top). Each colored line shows the tuning curve for a different neuron. With the Narrow prior, the tuning curves are narrower and more densely distributed around the prior mean.

We find that the variability of subjects’ estimates indeed increases with the width of the prior (Fig. 1B). With the Narrow prior, the across-subjects average standard deviation of estimates is about 4, while with the Wide prior (which is three times larger), it is around 7 (we estimate these quantities through Bayesian probabilistic methods; see Methods). A (pre-registered) Levene’s test rejects the equality of the variances across conditions (*p* < 0.0001; Bayesian methods yield the same conclusions; see Methods). More quantitatively, the variance of estimates increases linearly with the prior width, consistent with the results of the original experiment [10]. This linear scaling is predicted by an efficient-coding model formalizing a trade-off between the precision of estimates and the cost of this precision [10]. Our data thus suggest that in the context of numerosity representations, subjects implement fast efficient coding: they reorganize their representational resources on the order of one second, to adapt to changing stimuli statistics.

We note that a possible alternative account of the increase of the response variance with the prior width is that wider priors result in greater variability of Bayesian estimates. In other words, a Bayesian observer with unchanging encoding precision may also exhibit increasing response variability (see Ref. [23] for an example with a motion direction estimation task, in which subjects also exhibit increasing variability). Several arguments, however, support the hypothesis of a change in the encoding. First, an fMRI study of a very similar numerosity estimation task shows that the numerosity-encoding populations in human parietal cortex exhibit ‘distributed range adaptation’, i.e., their receptive fields shift and rescale when the range of the (uniform) prior changes [11]. Hence in this study the neural encoding does appear to change in adaptation to the prior. Second, we fit to the data several models of Bayesian estimation, including a model in which the internal, perceptual noise is insensitive to the width of the prior. We find that a large fraction of subjects is best fit by a model in which the variance of the internal noise scales linearly with the prior width, suggesting that their encoding adapted to the prior (more details on the models and the fits are available in Supplementary Information). For these reasons we further investigate adaptive modulations of the neuronal encoding, and how they may be mediated by gain adaptation in a recurrent sensory network.

### Dynamic tuning curves: Prior attraction and adapter repulsion

Traditional efficient-coding models posit a static prior and predict that the allocation of encoding resources should match this prior. Specifically, models of neural populations with unimodal tuning curves predict that the density of their locations should be proportional to the prior [3, 6–8] (or to the prior raised to some exponent, depending on the specifics of the model; here, for simplicity we consider the exponent 1). Integrating this relationship, one finds that a neuron’s ‘index’ (the integral of the density up to its location) corresponds to a given quantile of the prior (the integral of the prior up to the location). We are interested in the implications of ‘fast efficient coding’, i.e., the hypothesis that sensory networks quickly implement such optimal configuration, when the prior changes. To illustrate, we consider throughout this article the problem of encoding a scalar stimulus, *s*, drawn from a Gaussian prior with mean *µ* = 0, and a variable standard deviation, *σ*_*p*_. For the sake of illustration we consider three values, *σ*_*p*_ = 10, 20, or 30. (These widths match the parameters used in our estimation task, but the scalar stimulus considered here could represent any arbitrary one-dimensional variable, such as luminosity or loudness, and the specific scale chosen has no significant implications.)

If the prior narrows and efficient coding can be implemented quickly, then tuning curves should move closer to the prior center (Fig. 1D). More precisely, for the density of neurons to consistently match the prior in different contexts, each neuron’s location should correspond to the same quantile, across different priors (Fig. 1C). Note that here we have made the additional assumption that the neurons maintain their relative ordering, i.e., their ‘index’, across contexts. This assumption is consistent with empirical observations in various magnitude-coding systems (e.g., numerosity in humans [11], and time in rats [24]).

This ‘prior attraction’ of encoding resources would explain the changes in subjects’ precision in the numerosity estimation task. Indeed doubling the size of the prior would mean halving the density of neurons, i.e., each number would be encoded by half as many units as with the original prior, thus dividing the precision of the representation by two. (More precisely this would result in the estimate variance scaling with the square of the prior width; further below we comment on the reasons why the variance scales instead linearly with the prior width.)

Prior attraction is a prediction regarding the behavior of tuning curves when the relative frequencies of stimuli change. We note that the fMRI study mentioned above provides an empirical example of such prior attraction of encoding populations, in the context of a numerosity estimation task. But a number of experimental studies have documented a very different kind of change in sensory tuning curves following manipulations of the stimuli frequencies. In these experiments, typically, an adapter stimulus is repeatedly presented (Fig. 2A), and the common finding is that tuning curves shift *away* from this adapter (instead of toward the adapter as would be predicted by prior attraction) [13–18]. This ‘adapter repulsion’ is not a single phenomenon but rather encompasses a rich set of related effects observed in different systems. Specifically, we note three consistent experimental observations that have been made in different studies, in the primary visual cortices of macaque monkeys [14] and of cats [13, 15]. First, the tuning curves of sensory neurons whose locations are close to the adapter shift away from it. Second, this effect decreases with distance, and neurons with further locations shift less strongly. Third, neurons close to the adapter see their tuning curves widen, while those further away see their tuning curves become narrower. These stylized facts are illustrated in Figure 2B (solid and dashed lines). In addition, Refs. [13, 14] find some evidence for a slight attractive shift in neurons whose locations are even further away (Fig. 2B, dotted lines). In spite of this apparent adapter attraction, we label these observations taken together as ‘adapter repulsion’, since repulsion appears to be the dominant effect. Below, we examine a gain-adaptive recurrent-network model, and show how it accounts for both prior attraction and adapter repulsion under a single efficient-coding objective.

**Fig. 2:**
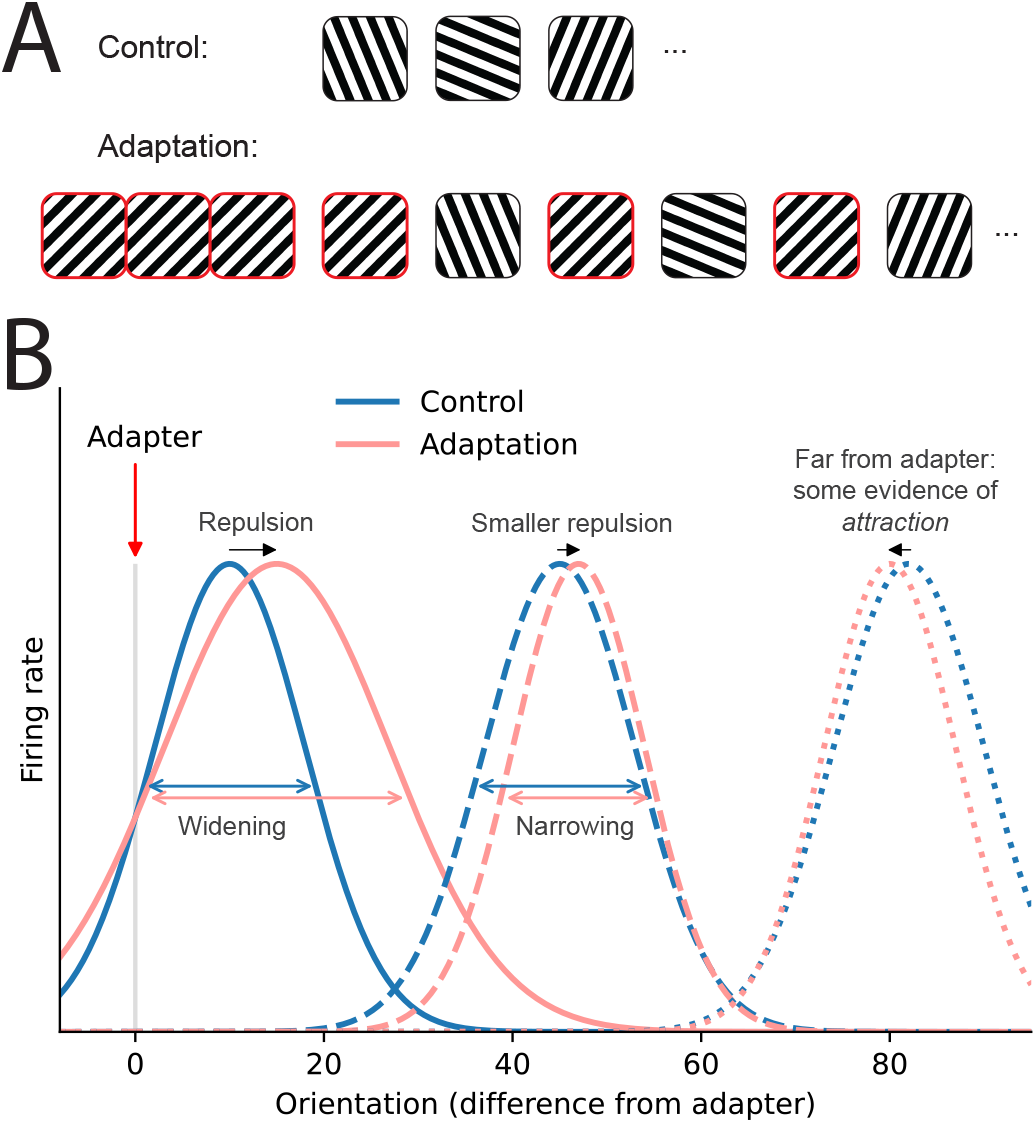
Adapter repulsion phenomena. **A**. Typical adaptation experiments (panel inspired by Dragoi *et al*., 2000 [13]). In the Control condition, a sequence of gratings with random orientations is shown. In the Adaptation condition, a single grating, the adapter, is presented for a long time or repeatedly. Then a sequence of gratings with random orientations, regularly interspersed with ‘top-up’ presentations of the adapter, is shown. **B**. Stereotypical results of adaptation experiments [13–15]. Tuning curves, i.e., firing rate as a function of the presented orientation, for three sensory neurons, in the Control condition (blue lines) and in the Adaptation condition (pink lines). For a neuron whose preferred orientation is close to the adapter (solid lines), the tuning curve widens and shifts away from the adapter. For a neuron whose preferred orientation is less close to the adapter (dashed lines), the tuning curve narrows and shifts away from the adapter, by a smaller extent. For a neuron whose preferred orientation is further from the adapter (dotted lines), some studies find a modest adapter attraction, i.e., the tuning curve slightly shifts towards the adapter [13, 14].

### Gain-adaptive recurrent neural network

We consider a network of *N* neurons encoding a stimulus *s* ∈ ℝ. Similarly to Ref. [22], weposit that their mean firing rates 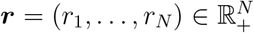 are determined, first, by the “feedforward tuning curves” response functions 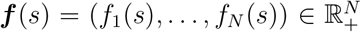, each multiplied by a gain, *g*_*i*_ ∈ ℝ_+_; second, by the recurrent activity of the network; and third, by random fluctuations. Specifically, we assume linear dynamics, as

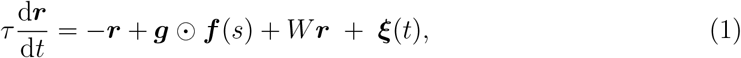

where ***g*** = (*g*_1_, …, *g*_*N*_) is the vector of the gains, ***g*** ⊙ ***f*** (*s*) is the Hadamard (elementwise) product of the vectors ***g*** and ***f*** (*s*), *W* ∈ ℝ^*N*×*N*^ represents the network’s net synaptic connectivity matrix (we do not model differentiated excitatory and inhibitory neurons), *τ* is a time constant, and ***ξ*** is a Gaussian white-noise term (𝔼 [***ξ***] = 0) which we detail further below. First, we are interested in the mean, steady-state activity of the network. Setting 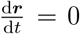 and taking the expectation yields the steady-state mean responses as functions of the stimulus *s*:

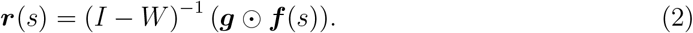

We call “effective tuning curves” the functions *r*_*i*_(*s*). They result from the feedforward tuning curves ***f*** (*s*), the gains ***g***, and the network’s recurrence *W*. Given a stimulus *s*, the effective tuning curves describe the neurons’ mean firing rates in the steady state.

We assume that the feedforward tuning curves *f*_*i*_(*s*) are Gaussian functions centered on the ‘feedforward locations’, *s*_*i*_, which are ordered and equally spaced by a distance *ℓ* > 0 such that *s*_*i*_ = *s*_1_ + (*i* − 1)*ℓ*. The curves have amplitude 1 and width parameter *σ*_*f*_, i.e.,

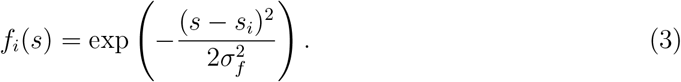

The connectivity matrix, *W*, is assumed to be structured such that the neurons whose locations are closer to each other are more strongly connected (this is consistent with studies on the connectivity of sensory networks, for instance in the mouse primary visual cortex [25– 27]). Specifically, we assume that the connectivity matrix is built with a Gaussian kernel with width parameter *σ*_*rec*_, as

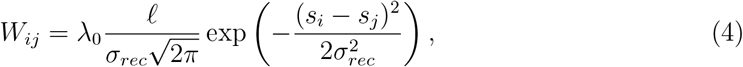

where 0 < *λ*_0_ < 1 is a parameter representing the overall strength of the connectivity (it is also an approximation of the largest eigenvalue of *W* ; see Methods). We also consider a model that includes general lateral inhibition, and obtain similar results (see Supplementary Information).

The random spikes emitted by the network’s neurons introduce random fluctuations in the dynamics of the rates, modeled in Eq. 1 by the stochastic term ***ξ***. Its covariance is thus determined by the variance of the Poisson spikes, which is equal to their rate: specifically, we approximate these random dynamics by positing 𝔼 [***ξ***(*t*)***ξ***(*t*′)^⊤^] = *τ* Γ*δ*(*t* −*t*′), where Γ = diag(***r***(*s*)). Below, we first have a closer look at the effective tuning curves, ***r***(*s*), before turning to the question of decoding the spike trains resulting from the fluctuating rates.

### Effective tuning curves

In the absence of recurrence (e.g., *λ*_0_ = 0), the effective tuning curves are simply the feedforward tuning curves, multiplied by the gains (i.e., ***r***(*s*) = ***g*** ⊙ ***f*** (*s*); Fig. 3A). With recurrence (*λ*_0_ *>* 0), the activity of each neuron is influenced by all the other neurons, to varying degrees that depend on ***g*** and *W*. With uniform gains (i.e., if all the gains *g*_*i*_ are equal), these lateral inputs impact the overall activity of the neurons, but not their locations (Fig. 3B). If instead the gains are non-uniform, then each neuron’s effective location will depend on the gain profile and on the recurrence matrix. Informally, a given neuron’s location is pulled towards the feedforward locations of the neurons with high gains, and this effect is stronger with nearby locations (Fig. 3C). Formally, Equation 2 provides the effective tuning curves, but not in a way that makes readily explicit their effective locations and widths. Here we provide analytical approximations to these quantities, that make them more transparent. Our derivations are detailed in Methods.

**Fig. 3:**
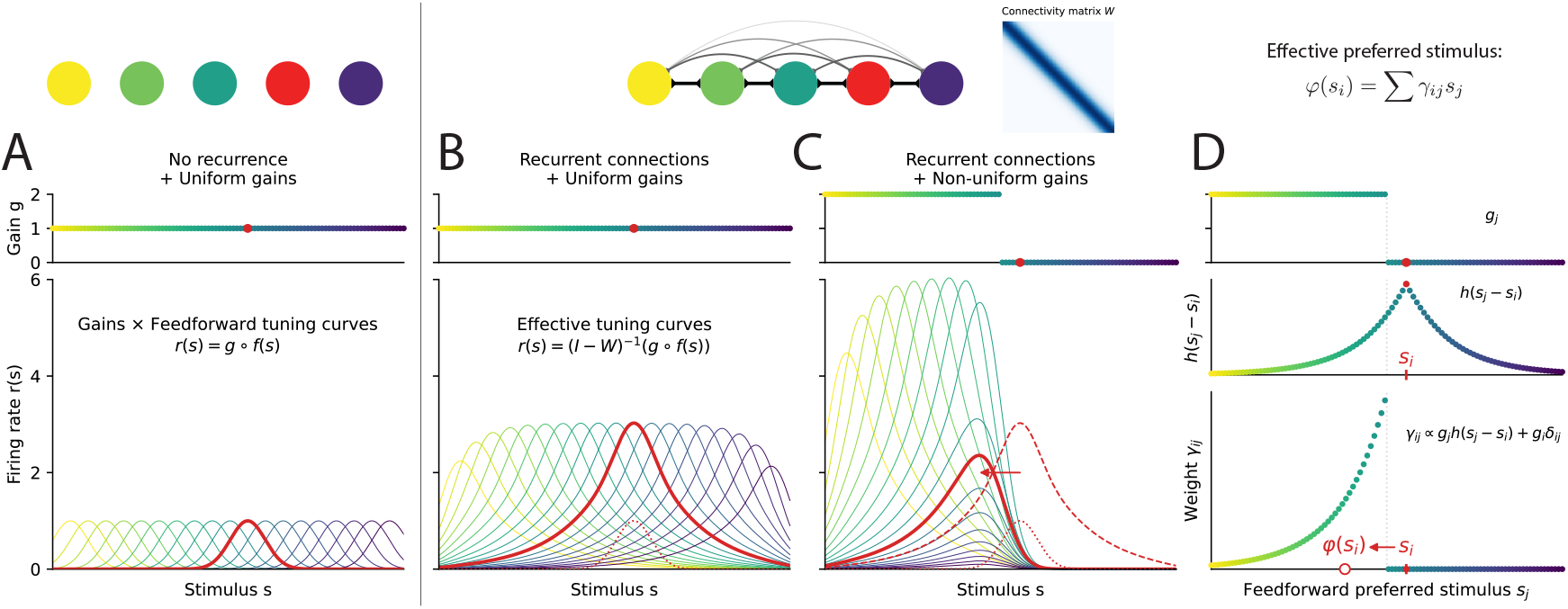
Tuning curve shifts via gain modulation in a recurrent network. **A**. Without recurrence and with gains uniformly equal to 1 (top), the effective tuning curves are not different from the feedforward tuning curves. Thus all the neurons have identical tuning curves, except for their locations (bottom). **B**. With recurrence, and maintaining the gains uniformly equal to 1, the activity of each neuron propagates in the network, resulting in a general increase of the amplitudes of the tuning curves, but no changes in their locations. The amplitude falls off near the edges of the stimulus space owing to the Toeplitz but non-circulant recurrence matrix. **C**. With non-uniform gains (here, half the neurons have gain 2 and half have gain 0), the tuning curves locations of the neurons with lower gain shift towards those of the neurons with higher gains. In red, tuning curve of a neuron (solid line), compared with its tuning curve with uniform gains (dashed line, solid line in **B**) and with no recurrence (dotted line, solid line in **A**). The inset shows the connectivity matrix (darker blue corresponds to stronger connections). **D**. The effective location of a neuron *i* is a weighted sum of the feedforward stimuli of all the neurons. The weights *γ*_*ij*_ (bottom panel) are determined by the gains *g*_*j*_ (top panel) and by a decreasing function *h*(*s*_*j*_ −*s*_i_) of the distance between the two neurons’ locations (middle panel).

We find that the effective location of neuron *i*, φ(*s*_*i*_), and its effective tuning-curve squared-width, 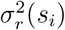 (defined respectively as the ‘average’ and the ‘variance’ of the tuning curves, see Methods), can be expressed approximately with weighted averages, as

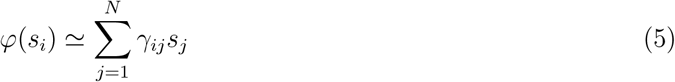

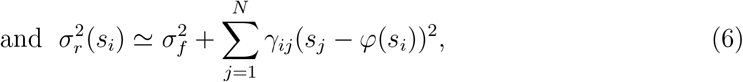

Where

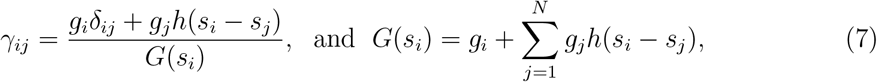

where *δ*_*ij*_ is the Kronecker delta, and *h*(*z*) is a decreasing function of |*z*| (see Fig. 3D, middle panel). The effective location of neuron *i*, φ(*s*_*i*_), is thus a weighted average of the feedforward locations *s*_*j*_ of all the neurons. The weight *γ*_*ij*_ of each neuron *j* is proportional to its gain, *g*_*j*_, and it decreases the further neuron *j* is from neuron *i*. In other words, neurons with greater gains “pull” towards them the other neurons, with a greater pull on the closer neurons. Figure 3D illustrates, for a neuron *i* (in red) and given a gain profile (top panel), the unimodal function *s*_*j*_ ↦ *h*(*s*_*j*_ − *s*_*i*_) (middle panel), the weights *γ*_*ij*_ of the neurons *j* (bottom panel), and the resulting effective location of neuron *i*, φ(*s*_*i*_), shown to be shifted away from its feedforward location, *s*_*i*_.

Finally, we use these results to approximate the neuron’s tuning curve as a Gaussian function. We obtain:

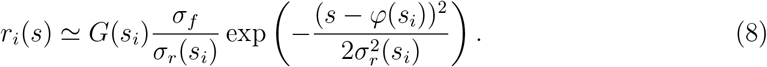

Equation 8 provides an analytical expression describing the probabilistic population code carried out by this network given a gain profile ***g***. The quality of our approximations of *W, φ*(*s*_*i*_), 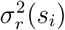, and *r*_*i*_(*s*) depends on the values of the various parameters, including *σ*_*f*_ (the width of the feedforward tuning curves), *σ*_*rec*_ (the width parameter of the connectivity matrix), *λ*_0_ (the strength of the connectivity), *ℓ* (the distance between two successive feedforward locations), and ***g*** (the gains). An important condition for the accuracy of our approximations is that the connectivity width, *σ*_*rec*_, should not be small compared to the neuronal spacing, *ℓ*. For the parameter regime used in this study (detailed below), the approximations are reasonably accurate (and many remain good even for *λ*_0_ = 0.99; see Supplementary Information for a comparison of these quantities with their approximations).

### Decoding error

What does the activity of this network tell us about the stimulus it is encoding? Considering, first, the case of fixed rates (without fluctuations), we look at the Bayesian posterior implied by the counts of spikes, ***k*** = (*k*_1_, …, *k*_*N*_), emitted by the network during one unit of time. As for the prior, we posit a Gaussian distribution, which we denote as 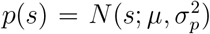. With our Gaussian approximation of the effective rates (Eq. 8), we obtain (see Methods) the approximate Gaussian posterior

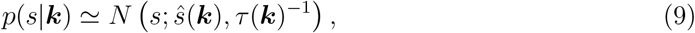

Where

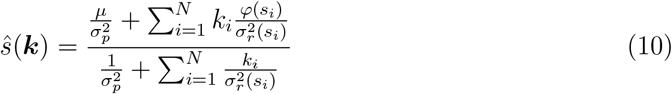

and the precision (inverse variance) is

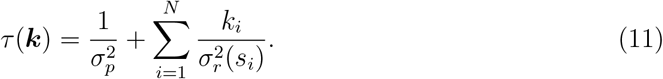

The optimal estimate, if one seeks to minimize the expected squared error, is the posterior mean, ŝ(***k***). Note that this estimator is a weighted average of the prior mean weighted by the prior precision, 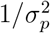, and of each neuron’s effective location, *φ*(*s*_*i*_), weighted by the number of spikes emitted by the neuron, *k*_*i*_, multiplied by its effective ‘precision’, 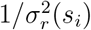. In other words, each spike is akin to a “vote” for the location of the corresponding neuron, weighted by a measure of the specificity of the neuron. This type of decoder has been known as population vector [28, 29], population average [30, 31], or center-of-gravity decoder [32]; see also Refs. [33, 34].

Furthermore, noting that the expected squared error of the Bayesian-mean estimator is the expected posterior variance, and using Jensen’s inequality, we find (see Methods) a useful lower bound on the mean squared error (MSE), as

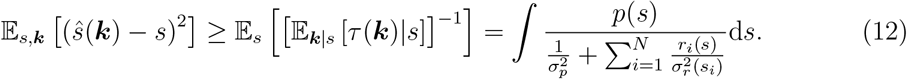

We note in addition that with the continuous approximations introduced below, this lower bound takes the perhaps more familiar form

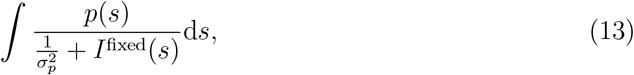

where *I*^fixed^(*s*) is the Fisher information of the network when the firing rates are fixed. As mentioned above, the rates are however subject to random fluctuations. These lower the information content of the network’s activity, as each spike stems from the combined stochasticity of the Poisson randomness and of the fluctuating rates. The resulting Fisher information of each spike, *I*_*i*_(*s*), is a fraction of the Fisher information that would obtain should the rates be fixed, i.e., 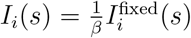, where *β* > 1 depends on the parameters of the network (see Methods). With our chosen parameters (detailed below), we have *β* ≈3*/*2, i.e., each spike carries two thirds of the information it would carry if the rates were fixed. In other words, the network with fluctuating rates is equivalent to a network with fixed rates, but with rates divided by 3*/*2. Therefore the lower bound on the MSE becomes

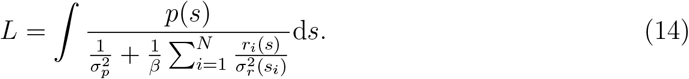

Below we use this lower bound *L* as our loss function. In our analyzes we have compared the squared error to the inverse of the expected precision, and we have found that the difference is relatively small, with our parameter regime (see Supplementary Information).

### Efficient coding under spiking cost

We now explicitly address the question of optimal coding. We consider an observer equipped with the described recurrent sensory network, and asked to estimate, under a quadratic loss, a presented stimulus *s* that is randomly drawn from the prior *p*(*s*). Prior to the presentation of *s*, we inform the observer of the prior *p*(*s*), as in our experiment above, and we allow for the adjustment of the gains, ***g*** = (*g*_1_, …, *g*_*N*_), of the sensory network. The synaptic connectivity matrix, *W*, and the parameters of the feedforward tuning curves, however, remain fixed.

Crucially, we posit that each spike is costly. In particular, we posit a cost proportional to the expected number of emitted spikes [35, 36], as

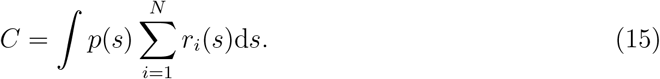

The gains are chosen so as to minimize the lower bound on the MSE derived above, *L*, under this cost, i.e., to solve the optimization problem

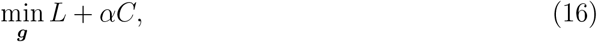

where *α* is a coefficient specifying the relative weight of the cost.

### Optimal gains under a prior larger than the tuning curves

Given a regime of parameters (i.e., the specifications of *σ*_*f*_, *σ*_*rec*_, *λ*_0_, *ℓ, N, σ*_*p*_, and *α*), the optimization problem just presented can be numerically solved, but it requires minimizing a function of the *N* parameters *g*_1_, …, *g*_*N*_, which can be computationally intensive. Here we provide instead an approximate optimal solution, which relies on the assumption that the prior is wide in comparison to the widths of the tuning curves (see Methods). We take the network of neurons to the continuous limit (*N* → ∞), and consider the functions *g*(*s*), *φ*(*s*), etc., instead of the discrete sets (*g*_1_, …, *g*_*N*_), (*φ*(*s*_1_), …, *φ*(*s*_*N*_)), etc., allowing us to approximate discrete sums with continuous integrals. An insightful intermediate result is

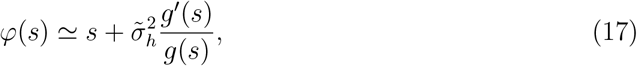

where 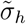 is proportional to *σ*_*rec*_ and depends on the other fixed parameters. Equation 17 shows that the shift of the effective location, *φ*(*s*), as compared to the feedforward location, *s*, is driven by the variations of the gains, *g*. Intuitively, the tuning curve locations are pulled towards neurons with greater gains (e.g., *φ*(*s*) *> s* if *g*′(*s*) *>* 0, and conversely if *g*′(*s*) < 0).

We solve the efficient-coding problem (Eq. 16) for the (continuous) gains *g*(*s*) by deriving the corresponding Euler-Lagrange equation under a small-fluctuations assumption on the gains, yielding an ordinary differential equation for *g*(*s*) wherever it is non-vanishing (*g*(*s*) *>* 0; see Methods). The approximate solution we obtain equals the positive part of an affine function of the log prior, as

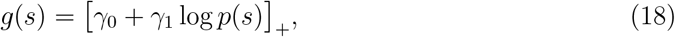

where *γ*_0_, *γ*_1_ are constants and [*x*]_+_ = max(0, *x*). Intuitively, the neurons whose locations are more probable under the prior should see their gains increased. This in turn determines the shift of the tuning curves, as the effective locations directly depend on the derivative of the gains (Eq. 17). Specifically, the gain profile in Eq. 18 implies that for the neurons whose feedforward locations are below the central value of the prior (*s* < *µ*), the effective locations will be greater than the feedforward one (*φ*(*s*) *> s*), while feedforward locations above the central value (*s > µ*) will result in lower effective locations (*φ*(*s*) < *s*). Put simply, the effective locations are attracted towards the central value of the prior, and thus the tuning curves contract towards the center.

### Gain adaptation yields prior attraction

We now optimize the gains of the recurrent network to three Gaussian priors, labeled ‘Narrow’, ‘Medium’, and ‘Wide’, with mean *µ* = 0 and widths *σ*_*p*_ = 10, 20, and 30, respectively (Fig. 4A). We look both at our approximation (Eq. 18) and at the numerically-optimized gains. For these analyzes we fix the various parameters as follows: *σ*_*f*_ = 5, *σ*_*rec*_ = 6, *λ*_0_ = 0.95, *ℓ* = 0.5, *N* = 801, and *α* = 0.5. These values are comparable to those chosen in similar studies, they yield tuning-curve widths of the same order of magnitude as those measured in numerosity-sensitive neurons [37–39], and result in estimate variability comparable to that obtained in the behavioral task. Finally, in our numerical optimization we add a regularization term on the second derivative of the gains, 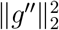, in order to avoid unstable solutions.

**Fig. 4:**
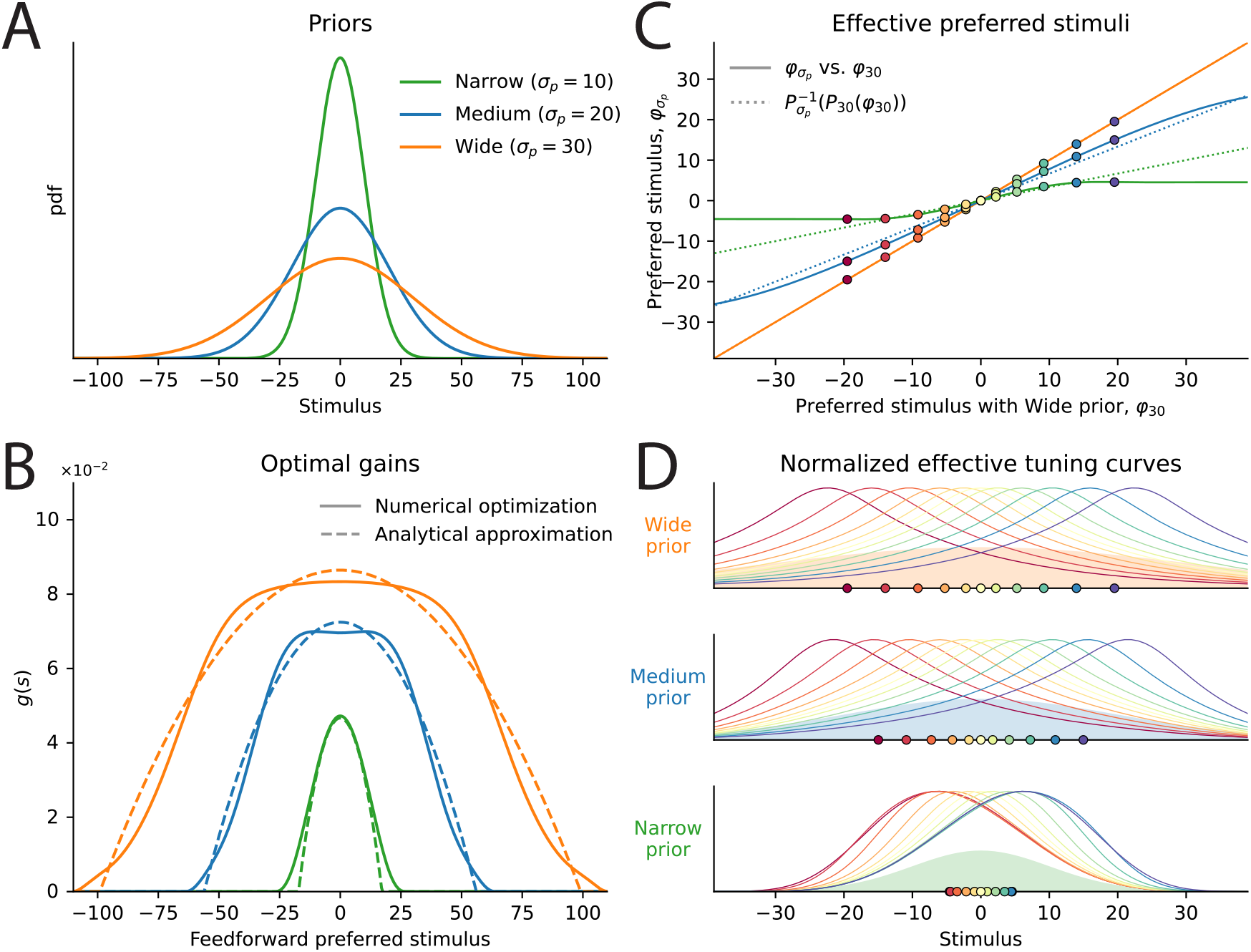
Optimal gains yield efficient coding through prior attraction. **A**. Narrow, Medium, and Wide Gaussian priors. **B**. Optimal gains *g*(*s*) as a function of the feedforward location, for the three priors, obtained through numerical optimization (solid lines) and approximate analytical derivation (Eq. 18; dashed lines). **C**. Neurons’ effective locations, 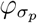, with the gains optimized to the different priors, as a function of their effective locations under the Wide prior, *φ*_30_ (compare to Fig. 1C). **D**. Normalized effective tuning curves, *r*_*i*_(*s*), exhibiting prior attraction resulting from the optimization of the gains (compare to Fig. 1D).

Our analytical approximation of the optimal gains is relatively close to the optimum we find numerically. The optimal gains have an inverted-U shape centered on *µ* (Fig. 4B); they are non-zero near the prior mean, and zero far from the prior mean: as a result, the effective locations of the network’s neurons are attracted towards the prior mean, and this contraction is stronger for the narrower priors (Fig. 4C,D). As mentioned above, fast efficient coding predicts that from one prior to the other, a neuron should encode for the same quantile. More precisely, denoting by 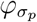 and 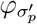 the effective locations of a given neuron under the priors with widths *σ*_*p*_ and 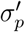, respectively, and by 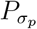 and 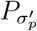 the CDFs of these priors, then the prediction is 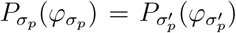. We thus look at the neurons’ effective locations under the Narrow and Medium priors, in comparison with those predicted on the basis of the Wide case, i.e., we compare 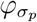 and 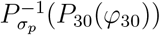 as a function of *φ*_30_, with *σ*_*p*_ ∈ {10, 20}. We find that the effective locations under the Medium prior are closer to the prior mean than under the Wide prior, and that they are well predicted by the relation just mentioned (Fig. 4C, blue lines). As for the locations under the Narrow prior, they are also well predicted by this relation, although only over an interval of roughly two standard deviations around the prior mean (Fig. 4C, green lines). As a result, the effective tuning curves exhibit ‘prior attraction’, i.e., they contract towards the center of the prior when the prior narrows (Fig. 4D). (Some obtained tuning curves have relatively long tails; we show in Supplementary Information that a model with inhibition suppresses the long tails.)

We emphasize that the mentioned prediction, 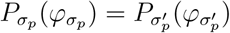, is obtained in efficient-coding models in which the locations are free parameters, optimized to an objective but otherwise unconstrained. By contrast, the efficient-coding problem we have considered is that of optimizing the gains in a strongly structured network of neurons. These gains together determine the effective locations, and we find them to behave similarly to the predictions of the “unconstrained” models, suggesting that recurrent networks can implement fast efficient coding through gain adaptation.

### Anatomy of the network’s Fisher information

The locations and shapes of the neurons’ tuning curves determine the precision of the encoding carried out by the network. We now examine a statistical measure of this precision, the Fisher information *I*(*s*). Generally, for a continuum of Poisson neurons with locations *s*, gain *g*(*s*), Gaussian tuning curves with width *σ*(*s*), and density *d*(*s*), if these functions varyslowly on the scale of *σ*(*s*), then the Fisher information is approximately (see Methods):

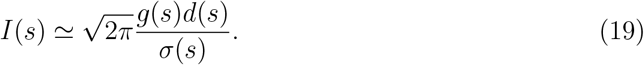

We examine, with our recurrent network, how these three quantities, *g*(*s*), *σ*(*s*), and *d*(*s*), vary with the prior width when the gains are adapted to the prior, and we look at how together they result in the Fisher information of the population. (Note that in our network the density is directly related to the locations, as 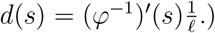. For simplicity, we only consider these quantities at the central value, *µ*.

The gains increase as a function of the prior width (Fig. 5A). This results in larger spiking activity for larger widths (i.e., the expected number of spikes increases; Fig. 5E, dashed line). As the prior widens, the tuning curves spread out, but they also widen, i.e., their density decreases and their widths increase (Fig. 5B). Informally, if tuning curves are half as dense, then they should double in width to cover the same ground. Here we find indeed that the effective widths of tuning curves are an affine function of the inverse of the density (Fig. 5C). The population’s Fisher information results from the combination of these three factors.

**Fig. 5:**
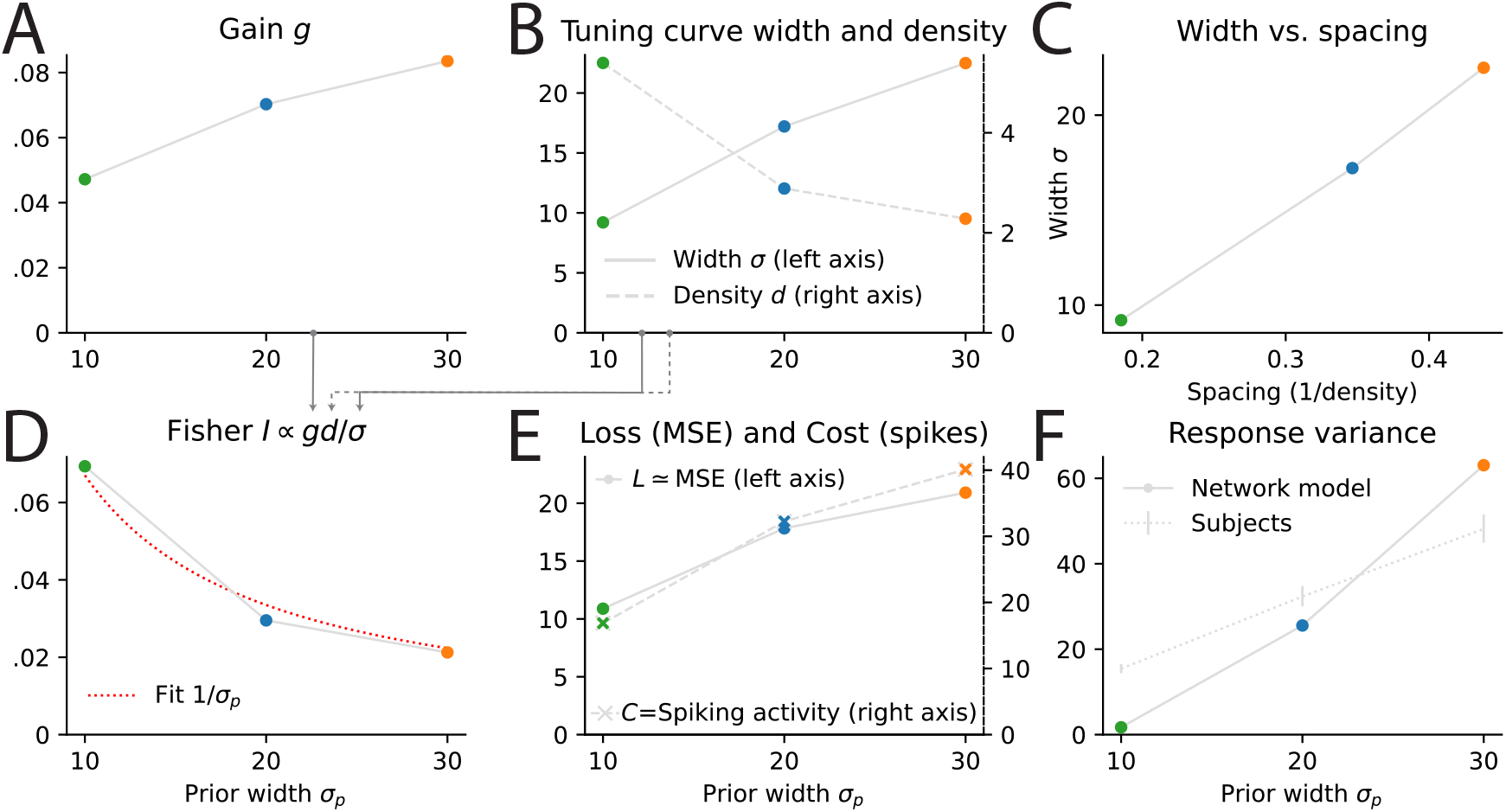
Properties of the encoding network across priors of different widths. Gain (**A**), tuning curve width and density (**B**), Fisher information (**D**), loss function *L* and cost *C* (**E**), and variance of simulated estimates (**F**), as a function of the width parameter of the Gaussian priors, *σ*_*p*_. **C**. Tuning curve width as a function of the inverse of the density (which indicates the spacing between tuning curves’ locations), exhibiting a linear relationship: as tuning curves become more widely spaced, their width increases. In **F**, the variance of subjects in the estimation task is also shown (dashed line; as in Fig. 1B).

We find that it decreases as a function of the prior width: informally, the neural population is less precise when there are more stimuli to be represented (Fig. 5D). The Fisher information as a function of the prior width, *σ*_*p*_, is well fit by a function proportional to 1*/σ*_*p*_. As a result, the variance of the estimates derived from the population’s simulated activity is an increasing function of the prior width (Fig. 5F), similarly to subjects’ behavior (Fig. 1B and Fig. 5F, dashed line). We also look at the standard deviation of the posterior, as a measure of the observer’s uncertainty, and we find that it increases with the prior width (see Supplementary Information).

The dependence of the Fisher information in 1*/σ*_*p*_ (rather than 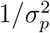) and of the variance in *σ*_*p*_ (rather than 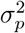) reflects the endogenous choice in precision that results from the efficient-coding problem that features a cost on the spiking activity. The terms of the trade-off, *L* and *C* (Eqs. 14-15), are shown in Figure 5E. The loss, *L*, is the expected mean squared error (MSE). If the network dedicated the same amount of encoding resources in the context of the different priors, the loss would increase as the square of the width, 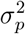. This would result in large losses. But the MSE in fact increases at a lower rate (Fig. 5E, solid line), because the gains increase with larger priors (Figs. 4B and 5A), resulting in an intensification of the spiking activity (Fig. 5E, dashed line). This increased activity is costly, but it optimally mitigates the worsening of the loss [10].

### Peaked prior and adapter repulsion

The analytical results above rely on approximations that are less accurate when the prior width is close to, or lower than, the widths of the tuning curves. What are the optimal tuning curves under very narrow, that is, “peaked” priors? We consider this question in relation with the experimental finding that the tuning curves of some sensory neurons, after repeated exposure to an ‘adapter’ stimulus, shift away from this stimulus, as detailed above (Fig. 2). Hence we examine the tuning curves of the recurrent network in a scenario similar to these studies. It is not clear what the prior should be after repeated presentations of a stimulus; thus to simplify we consider a ‘control’ prior (pre-adaptation) that is uniform over a large interval, in comparison with a peaked, ‘adaptation’ prior that is a mixture between the same uniform distribution (for 80%) and a very narrow Gaussian prior of width *σ*_*p*_ = 1 and centered on *µ* = 0 (for 20%; Fig. 6A; see [4] for a similar approach).

**Fig. 6:**
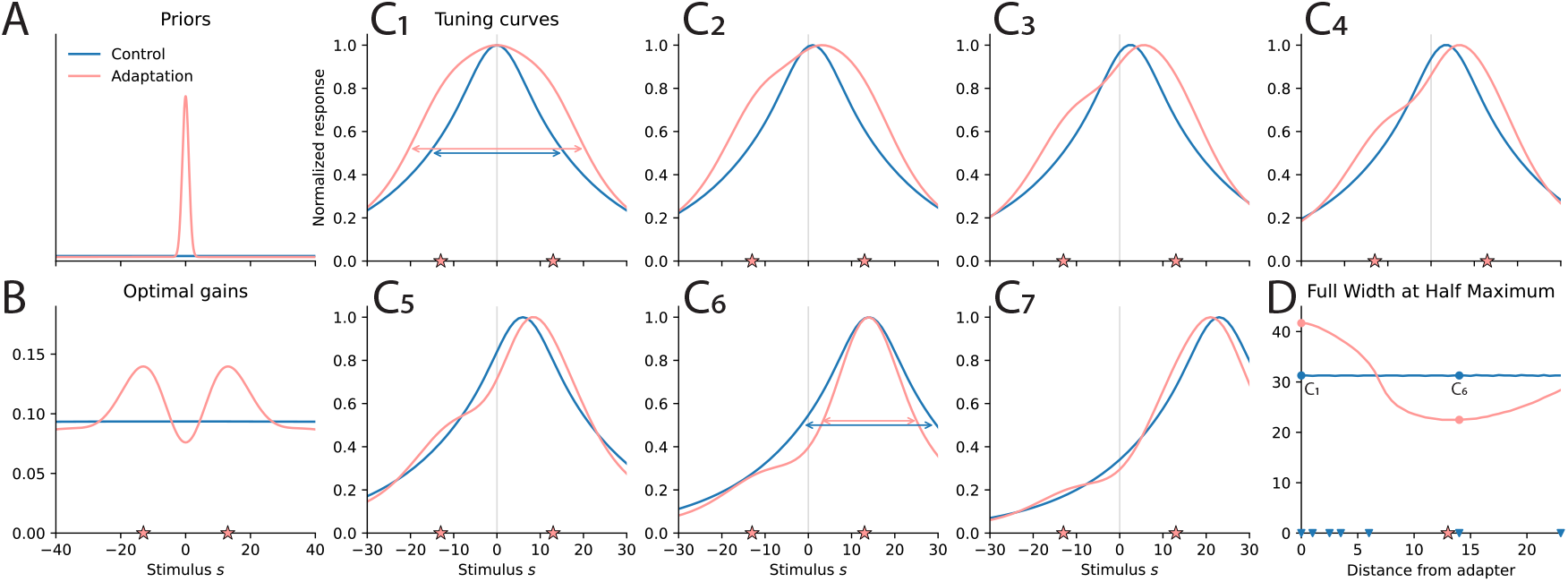
Efficient coding via gain adaptation captures the rich adapter-repulsion phenomena. **A**. Uniform, Control-condition prior (blue line), and Adaptation-condition prior (pink line). **B**. Gains optimized for the two priors. With the Adaptation prior, the optimal gains exhibit two local maxima whose locations, marked with pink stars, are away from the peak of the prior. **C1-7**. Tuning curves for seven different neurons, in the two conditions. The pink stars indicate the locations of the gains’ maxima. In the Adaptation condition, the tuning curves are ‘attracted’ towards these maxima, resulting in changes in their locations and widths. In C1 and C6, the widths of the tuning curves are shown, to emphasize the widening (C1) or the narrowing (C6). **D**. Full Width at Half Maximum of the neuronal tuning curves as a function of the distance from the adapter of their locations, in the two conditions. On the abscissa, the triangles show the locations of the neurons depicted in C1-7, while the pink star shows the location of the gains’ maximum. The dots on the curves corresponds to the neurons whose widths have been emphasized in C1 and C6.

The control prior provides the baseline tuning curves relative to which ‘attraction’ and ‘repulsion’ under the peaked prior are defined. We numerically optimize the gains to these priors. Importantly, we keep all the other parameters of the network fixed and identical to the values that we have used above. Hence it is exactly the same efficient-coding optimization problem (Eq. 16) that is solved: only the prior changes.

With the peaked prior, the optimal gains exhibit two local maxima symmetrically located away from the peak, while at the peak (*µ* = 0) the gains reach a local minimum, resulting in a bimodal M shape of the optimal gains as a function of the location (Fig. 6B). (We also vary the width of the Gaussian peak and the proportion of the uniform distribution in the adaptation prior, and we find that this M shape is robust to these changes; see Supplementary Information.) The shape of the gains in turn determines the shapes of the neurons’ tuning curves. For a neuron whose location is exactly at the adaptation peak, the tuning curve is “pulled” by the two gain maxima in both directions, resulting in a tuning curve almost twice as large as that obtained under the control prior (Fig. 6C1).

The neurons not located at the peak are unequally impacted by the two gain maxima, resulting in asymmetric tuning curves. The maximum of each curve is located, roughly, between the neuron’s location under the control prior (which is essentially the feedforward location), and the location of the closest gains’ maximum (Fig. 6C2-7). Thus for locations between the adaptation peak and the gains’ maximum, there is a repulsive shift (away from the peak; Fig. 6C2-5). In other words, the adapter repulsion, in this model, is understood as an attraction towards the gains’ local maximum. Feedforward locations further away from the peak and closer to the gains’ maximum result in shorter shifts, and the shift vanishes for a location close to the gains’ maximum (Fig. 6C6). Finally, for locations beyond the gains’ maximum, we obtain a shift towards the peak, i.e., ‘adapter attraction’, but it is better understood here as an attraction towards the gain maximum (Fig. 6C7).

The widths of the tuning curves, similarly, are determined by the bimodal gain profile. The ‘Full Width at Half Maximum’ (FWHM) is maximal for neurons whose locations are at the adaptation peak (Fig. 6C1). Further, the impact of one maximum decreases while that of the other increases, resulting in narrowing tuning curves (even narrower than under the control prior; Fig. 6C2-7). The minimum FWHM is reached near the location where the gains reach their local maximum (Fig. 6D). In sum, our model captures all the experimental findings that we have found to be consistently reported in the literature (and illustrated in Fig. 2B): namely, repulsive shifts close to the adapter, weaker repulsive shifts further from the adapter, and attractive shifts even further; also the widening of the tuning curves close to the adapter, and their narrowing away from it [13–15].

The sizes of the effects shown in Figure 6 are of the same order of magnitudes as those reported in the literature. Roughly, the repulsive shifts reported in Refs. [13–15] range from about 5% to 20% of the tuning curve widths (that is, 2.5º to 10º vs. widths of about 50º), with attractive shifts of about -5%. We obtain comparable effects, with repulsive shifts of up to 13% (Fig. 6C4), and attractive shifts of up to -6% (Fig. 6C7; here a shift is defined as the difference between peak locations, divided by the FWHM in the control condition). As for the changes in width, Felsen *et al*. (2002) report tuning curves widening by an average of 15% near the adapter, and narrowing by about -5% further from the adapter [15]. In our simulations we obtain a 33% widening (ratio of FWHMs) for the neuron whose location is exactly at the adapter (Fig. 6C1), and a narrowing of up to -28% (Fig. 6C6), i.e., stronger effects than in the data; we note that when introducing inhibition in the network, we obtain more moderate effects (see Supplementary Information). The parameters of the model were not fit to any experimental data, but these results show that the gain-adaptive network is able to produce tuning curve shifts and width changes that are quantitatively comparable to those observed experimentally. We note that, regarding the behavioral data, the same model yields a relationship between the variance of estimates and the prior width that has a slope comparable to that of subjects (Fig. 5F). This simultaneous quantitative agreement across very different empirical phenomena (and obtained with the same network’s parameters in both contexts) supports the gain-adaptive network model as a coherent conceptual framework.

## Discussion

We have studied a computational model of sensory representations that predicts a diverse range of adaptive changes in neuronal tuning curves following changes in stimulus frequencies, under a single principle — efficient coding — and using a single mechanism — neuronal gain adaptation. We presented behavioral evidence obtained in a pre-registered estimation task suggesting that human observers dynamically adjust their perceptual encoding in response to rapidly changing priors. Motivated by these findings, we introduced a gain-adaptive recurrent neural network in which neuronal gains minimize an objective combining decoding error and a metabolic cost proportional to neuronal spiking. Using analytical approximations and numerical simulations, we showed that this optimization yields prior attraction under wide stimulus priors and adapter repulsion under sharply peaked stimulus distributions, thus unifying apparently contradictory adaptive patterns. Our model suggests that fast efficient coding and the rich phenomena of adapter repulsion can be obtained through gain modulation alone, without requiring changes in synaptic connectivity.

Our behavioral experiment builds on previous work that had demonstrated subjects’ adaptation to the prior when it is manipulated during the experiment but unchanging in blocks of successive trials [10], a finding substantiated by behavioral results obtained with similar experimental settings [12, 40–42], and by fMRI data exhibiting distributed range adaptation of numerosity-sensitive populations in parietal cortex [11]. Ni and Stocker (2023) [41] in addition re-analyze data from visual averaging tasks and argue that efficient adaptation occurs at each trial. Our experiment explicitly tests this hypothesis by examining behavior when the prior distribution changes from one trial to the next. Our results replicate those obtained previously with blocked designs [10], supporting the hypothesis that sensory representations efficiently adapt to changing priors on rapid timescales (on the order of one second).

Several neurophysiological studies support the idea that neural responses dynamically adapt to changing stimulus statistics in a manner consistent with fast efficient coding [43, 44]. In the blowfly visual system, H1 neurons (selectively responsive to horizontal motion) adjust their responses when the variance of the stimulus distribution changes [45, 46]. Specifically, the (monotonic) tuning curves are invariant when plotted as a function of the velocity normalized by the standard deviation of the prior. Similarly, in cat lateral geniculate nucleus [47], in songbird auditory neurons [48], and in rat barrel cortex [49], gains have been showed to vary as a function of the variance of the stimulus amplitude.

Related observations have been made in the domain of value-based decision-making: dopamine neurons adapt their coding to the range of reward magnitudes [50], and neurons in the orbitofrontal cortex dynamically adjust their response functions to represent economic value across different ranges [51]. Most of these examples involve monotonic tuning functions. Two studies in which tuning curves seem to be unimodal functions of the stimulus report results consistent with fast efficient coding. These studies investigate the encoding of time in the rat hippocampus [52] and striatum [24]. Both works report a scalable representation of time, with patterns of neural activity that stretch or compress in response to changes in the duration being timed. This suggests a scalable and dynamic encoding — although not on a trial-to-trial basis, in these studies, but on the timescale of experimental sessions.

Fast efficient coding, in our model, is mediated by gain modulation. A substantial body of research has explored the role of gain modulation in shaping neuronal responses, and in particular how it may benefit coding efficiency. Gain modulation has been proposed as a multiplicative mechanism by which an ‘attention field’ is implemented, in order to improve performance in a given task [53]. Gain control also enables the decorrelation of multidimensional inputs, thereby carrying out a form of efficient coding that eliminates statistical dependencies in natural signals [54, 55]. The adaptations of gains to the variance of stimuli, mentioned just above [47–49], are examples of fast efficient coding mediated by gains, in single neurons. They are however specific to monotonic tuning curves, as in these cases gain adaptation essentially allows for mapping the range of stimuli onto a limited range of firing rates. The role of gains in the mechanism of fast efficient coding that we have investigated here is of a different nature. Indeed here the unequal gains across the recurrent population induce shifts in the neurons’ locations, which, when optimized to a prior, allow for an efficient ‘coverage’ of the stimulus space. Hence this mechanism inherently operates at the population level (it does not apply to single neurons), and it is intrinsically tied to the recurrent nature of the considered population.

Some models of adapter repulsion also leverage the recurrence of sensory networks, but through synaptic plasticity. Westrick *et al*. (2016) proposes a divisive-normalization model of adaptation in which the normalization weights between each pair of neurons are updated through a Hebbian rule [20]. Such synaptic adaptation implies updating a number of parameters that scales as the square of the size of the neural population, which may yield instability and overfitting [22]. Perhaps more importantly, these synaptic updates typically occur on timescales longer than those of the adaptation phenomena we have considered here—on the order of a second, maybe less, for fast efficient coding, and less than 100ms for adapter repulsion.

A faster synaptic mechanism, often mentioned in studies reporting adapter repulsion [13– 15], is short-term synaptic depression, whereby the amplitude of postsynaptic responses is scaled down whenever the presynaptic neuron fires an action potential [19]. This synaptic-level gain control adapts the sensitivity of postsynaptic neurons to the fluctuations in the firing rates of afferent neurons. Such short-term plasticity occurs on a short timescale, but its effects spontaneously relax within hundreds of milliseconds. This limits its suitability for efficient coding, where the optimal gain configuration should be maintained as long as need be. Similarly, Quiroga *et al*. (2016) [21] show that repulsive shifts can emerge from the transient dynamics of a recurrent network, an account that does not resort to synaptic plasticity, but that predicts only short-lived adaptive effects.

The alternative mechanism investigated here is that variations in the neuronal gains of a recurrent circuit induce shifts in the location of tuning curves. Dragoi *et al*. (2000,2002) allude to this possibility [13, 14], and Duong *et al*. (2023) show that, under an efficient-coding objective, it predicts tuning shifts away from an adapter [22]. We build on this study and significantly extend it in several key ways. We provide analytical approximations to the main parameters of the effective tuning curves, as well as the optimal gains under wide priors, thus providing clear insights into the otherwise opaque behavior of the network; we introduce (and solve) an efficient-coding problem that includes only two terms, both readily justified: the loss function is derived from optimal, Bayesian inference, and the cost is proportional to the total expected number of action potentials, each of which consumes some amount of energy [35, 36]; we show in addition how our model captures the subtle features of adapter repulsion phenomena, including the widening and narrowing of tuning curves and the attraction far from the adapter; simultaneously within the same framework we predict prior attraction (and provide supporting behavioral data), thereby tying under a single paradigm the literature on efficient coding—traditionally understood as an optimal adaptation to a static prior—and that on adapter repulsion—the dynamic adaptation to a repeated stimulus.

The idea that adapter repulsion results from a form of efficient coding has been proposed elsewhere [4, 15, 16, 22], although a 2014 review on sensory adaptation concludes that efficient coding is “unlikely to be the sole goal of adaptation” [56]. Our results support the hypothesis that efficient coding is indeed the cause (if not the goal) of some adaptive phenomena, but our findings also suggest that this might be obscured by the non-trivial implications of efficient coding. Indeed efficient coding sometimes yield counter-intuitive predictions, for instance ‘anti-Bayesian’ percepts wherein perceptual estimates are pushed away from the more frequent stimuli [9, 57]. Here, encoding tuning curves are pushed away from the adapter. More precisely, we have seen that with some priors the tuning curves were attracted towards the prior, while with narrower priors they are repelled from it. Further, we obtain adapter repulsion and adapter attraction simultaneously, depending on the distance between the location and the adapter (as experimentally observed). Our model explains this seemingly contradictory behavior: in both cases tuning curves are attracted to a local peak of the gains; and the gains exhibit such peaks because they are optimally adapted to the statistics of the stimuli.

We have focused on the steady-state of the network, as in many studies on adaptation, although there are rich transient dynamics during adaptation [14, 18, 21]. We conjecture that gain changes first have an impact “locally” (i.e., on the neurons whose gains have changed), before propagating throughout the network. Thus there should be some latency in the tuning shifts, and the tuning curves of the neurons whose locations are further from the adapter should adapt later than the closer ones. This roughly corresponds to the findings reported by Dragoi *et al*. (2002) [14]; but we leave to future studies the investigation of the recurrent network’s transient dynamics.

Although our focus is on adaptive changes in sensory encoding, an important assumption of our model concerns how downstream circuits read out the adapted code. Throughout, we assume that decoding remains approximately optimal given the current encoding, in the sense that downstream neurons have access to (or learn on a fast timescale) the effective tuning curves induced by gain adaptation. Under this assumption, changes in encoding precision (e.g., a decrease in Fisher information, as in Fig. 5D) necessarily translate into changes in estimation precision (e.g., an increase in response variance, as in Fig. 5F). An alternative class of models assumes that encoding adapts while decoding remains fixed; in these models additional behavioral effects can arise from a mismatch between the decoder and the adapted code [58]. In our setting, however, maintaining a fixed decoder while the encoding reorganizes would generally lead to severely distorted estimates, making it difficult to interpret the resulting behavior. We thus focus on the case of optimal decoding and leave open the question of how downstream circuits might adapt their readout in parallel with changes in sensory encoding.

We have treated changes in stimulus statistics in the same way regardless of whether the corresponding priors are explicitly instructed, as in our numerosity-estimation task, or implicitly learned through repeated exposure, as in sensory adaptation paradigms. These two scenarios may differ in their learning dynamics. In an inference task, Berniker *et al*. (2010) demonstrate that human observers are able to learn both the mean and the variance of a prior, with distinct learning timescales [59]. While our numerosity-estimation task relies on explicitly cued priors, Heng *et al*. (2020) show that in a numerosity-discrimination task observers seem to adapt their encoding to different power-law distributions of numerosities [60]. In the domain of color perception, Eissa and Kilpatrick (2023) show that humans exploit learned environmental priors in a delayed estimation task [61].

Hence we have assumed that in adaptation experiments, sensory encoding is adjusted to the recent empirical distribution of stimuli. The implicit assumption is that the distribution of stimuli expected in the near future is approximated by the distribution observed in the recent stimulus history. Effectively, in these experiments, this distribution is strongly peaked around the adapter stimulus. A similar approach is adopted by Wainwright (1999) — also in the context of modeling visual adaptation as resulting from a form of efficient coding [4]. An implication of this perspective is that adaptation phenomena are inherently tied to the problem of online prior learning. The question of distribution learning, however, has received relatively limited attention. Among the studies mentioned above, Berniker *et al*. (2010) formalize prior learning as Bayesian inference over the parameters of a prior distribution [59], whereas Eissa and Kilpatrick (2023) consider a circuit-level model in which learned environmental priors are reflected in the formation of local attractor dynamics, mediated by synaptic plasticity [61]. The gain-adaptation mechanism we study here suggests a different route that does not resort to synaptic connectivity. Here we do not, however, model the history-dependent process of learning; how the gain changes are themselves learned or regulated over sequences of stimuli remains an important direction for future work.

The question of the dynamic adaptation of sensory systems to changes in stimuli statistics calls for considering the frequency of such changes, their ecological relevance, and the cost of adaptation. The frequencies of changes in environmental statistics cover several ranges of magnitudes, and depend on the timescale at which these statistics are considered. The change in luminance distribution when going from a shaded area to direct sunlight occurs at a relatively low frequency (but it is presumably important for an organism to quickly adapt to the new environment). By contrast, the local statistics of the visual patch covered by the receptive field of a primate’s V1 neuron change every time the eye makes a saccade, i.e., several times per second. Dragoi *et al*. (2002) report tuning curve adaptation over timescales of this order, in awake, behaving macaque monkeys, and suggest that it improves overall feature discrimination [14]. The mechanism of gain adaptation, however, probably comes with a cost (e.g., neuromodulatory signals), although we note that this cost is presumably lower than that entailed by synaptic reconfiguration, as suggested by other accounts of sensory adaptation. We have modeled, essentially, the efficient adaptation to single changes in stimuli statistics, neglecting the cost of gain adaptation. An interesting avenue for future research would be to examine a more general optimization problem, that would take into account not only the frequency of changes, but also the ecological benefit of sensory adaptation and the metabolic cost of gain adaptation.

In this study we have not modeled separate excitatory and inhibitory populations, and the matrix *W* should be interpreted as a net effective connectivity summarizing the recurrent interactions. A more biophysically complete model would explicitly distinguish excitatory and inhibitory neurons and specify the circuit mechanisms (e.g., normalization, disinhibitory control) through which such effective connectivity arises. Another prominent feature of biological neural circuits is homeostasis, the regulation of population activity to prevent runaway excitation and maintain stable operating regimes. It is possible that homeostatic mechanisms could induce changes in tuning curves, including stimulus-dependent reweighting of neural responses. An alternative hypothesis is that efficient coding gives rise to homeostatic properties, rather than the other way around. Indeed, first, the spiking cost penalizes excessive activity, thereby preventing firing rates from diverging. Second, gain-dependent shifts of tuning curves redistribute the encoding resources across stimulus space, ensuring that a large fraction of neurons remains engaged in the representation as stimulus statistics change. Together, these effects tend to stabilize population activity across contexts while preserving sensitivity to stimulus statistics. This interpretation is consistent with recent work showing that efficient-coding principles can simultaneously explain neural response homeostasis and stimulus-specific adaptation [62].

## Methods

### Effective tuning curves

In order to obtain an analytical approximation of ***r***(*s*), we derive first an approximation of *M* = (*I* −*W*)^−1^ through the diagonalization of *W*. The eigenfunctions of a continuous Gaussian kernel can be expressed as a combination of cosine functions; in the discrete case, the ‘discrete cosine transform’ provides a way to obtain approximate eigenvalues and eigenvectors of the matrix *W*. From these we can derive their counterparts for *M*. This enables us to find an approximate expression for the elements *M*_*ij*_, as *M*_*ij*_ ≃ *δ*_*ij*_ + *h*(*s*_*i*_ − *s*_*j*_), where *δ*_*ij*_ is the Kronecker delta, and *h*(*z*) is a decreasing function of |*z*| (see Fig. 3D, middle panel). Depending on the strength of connectivity, *λ*_0_, the function *h* resembles either a Gaussian function (small *λ*_0_) or a Laplace function (large *λ*_0_; i.e., a function proportional to exp(−|*z*|*/*ν), for some width ν *>* 0). The latter has heavier tails, implying a greater range of each neuron’s influence. The details of the mathematical derivations can be found in Supplementary Information.

Substituting *M* in Eq. 2 with its approximation yields an approximation of the effective tuning curve of a neuron *i, r*_*i*_(*s*), as a weighted sum of the feedforward tuning curves of all the neurons:

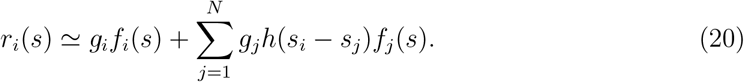

Thus the contribution of neuron *j* to the effective tuning curve of neuron *i* ≠ *j* is equal to its feedforward tuning curve *f*_*j*_(*s*) multiplied by its gain *g*_*j*_ and by a decreasing function of the distance between the feedforward locations of the two neurons, *h*(*s*_*i*_ − *s*_*j*_).

We then define the effective location of neuron *i, φ*(*s*_*i*_), and its effective tuning curve squared-width, 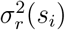, as

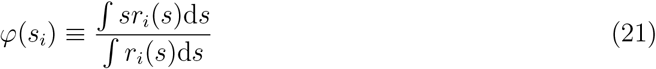

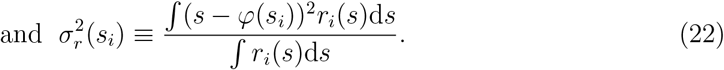

With the approximation of *r*_*i*_(*s*) in Eq. 20, we obtain expressions for *φ*(*s*_*i*_) and 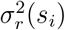 (Eqs. 5,6).

### Bayesian decoding with fixed rates

We first consider Bayesian decoding in the absence of rate fluctuations, i.e., with independent, Poisson-distributed spikes with fixed rates *r*_*i*_(*s*).

#### Posterior

We examine the Bayesian posterior implied by the spikes emitted by the network during one unit of time. We denote by ***k*** = (*k*_1_, …, *k*_*N*_) the vector of the counts of spikes emitted by the *N* neurons. The likelihood *p*(***k***|*s*) is

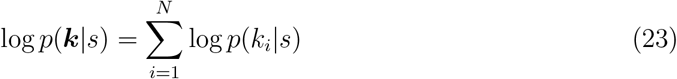

Where

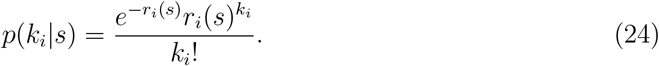

The Fisher information of neuron *i*, in this fixed-rate case, is thus

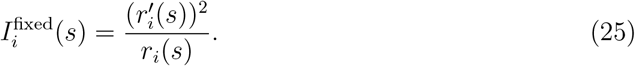

The log-posterior is

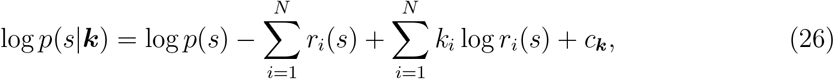

where *c*_***k***_ is a constant that depends on ***k*** but not on *s*. As for the prior, we posit a Gaussian distribution, which we denote as 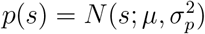. As for the sum of the rates, 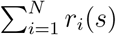, we assume that it varies slowly on the scale of the width of the posterior and thus that it can be neglected here—we note that in many similar models, this term is simply assumed constant [34, 63, 64] (we also further justify our assumption in Supplementary Information). With our Gaussian approximation of the effective rates (Eq. 8), we obtain the approximate Gaussian posterior presented in Eq. 9.

#### A lower bound on the Bayesian MSE

The unconditional, expected squared error of the Bayesian-mean estimator is the expected posterior variance, as

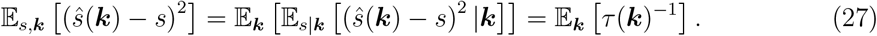

In turn, the unconditional expected posterior variance is naturally expressed (by the law of total expectation) as

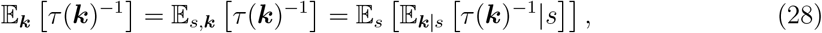

where 𝔼_***k***|*s*_ [*τ* (***k***)^−1^ *s*] is the expected value, conditional on a stimulus *s*, of the posterior variance. By Jensen’s inequality, a lower bound on this quantity is the inverse of the expected posterior precision, conditional on *s*, i.e.,

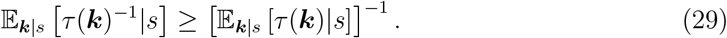

This conditional expected precision is readily derived from Eq. 11 as

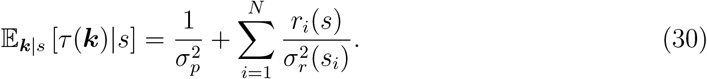

Thus a lower bound on the mean squared error, unconditional on *s*, is the expected value, taken over *s*, of the inverse of the conditional expected precision, which we denote by *L*^fixed^, i.e.,

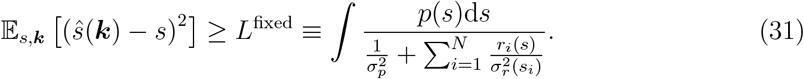

### Continuous approximations

Because we consider the case of relatively wide priors, we assume that the various functions characterizing the network (*g*(*s*), *φ*(*s*), etc.) vary slowly on the scale of the neurons’ widths. In addition to Eq. 17, we obtain the following useful continuous approximations (see Supplementary Information). The effective squared-width is approximately

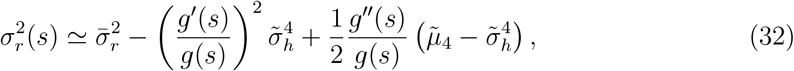

where 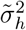 is a constant that represents the squared-width of the function *h*, and that is proportional to 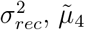 is a ‘fourth-moment’ quantity characterizing *h* (it is proportional to 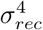); and where we denote by 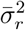 the typical width of the tuning curves:

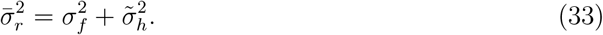

This is in particular the width of the tuning curves when the gains are uniform (*g*′ = 0). The total expected number of spikes, given a stimulus *s*, is approximately

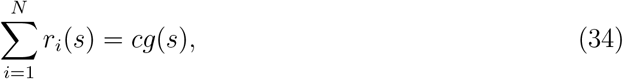

Where

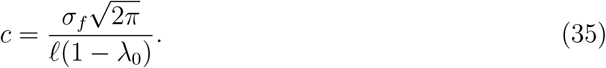

In other words, the expected number of spikes emitted by the whole network, conditional on a stimulus *s*, is essentially determined by the gain of the neuron whose location is *s*. Finally, the expected precision of the decoder (Eq. 30) is driven by the quantity

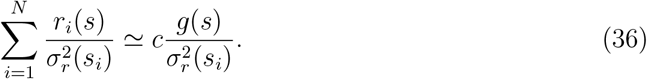

This is also the approximate Fisher information, i.e.,

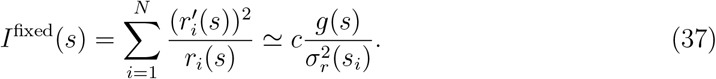

Therefore the lower bound on the expected loss is written as

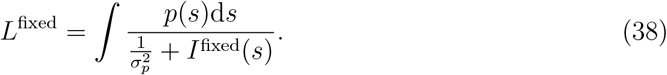

### Rates fluctuations

The fluctuations, *δ****r***, of the firing rates around their mean, ***r***(*s*), verify the dynamics equation

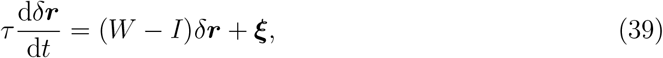

where 𝔼 [***ξ***] = 0 and 𝔼 [***ξ***(*t*)***ξ***(*t*′)^⊤^] = *τ* Γ*δ*(*t −t*′). At the equilibrium, the steady-state covariance matrix of the rates, Σ = 𝔼 [*δ****r****δ****r***^⊤^], verifies the Lyapunov equation

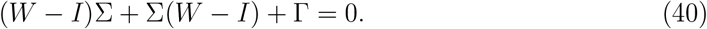

The stochasticity in the network stems from the random Poisson spikes emitted by the neurons, thus we posit that the instantaneous noise covariance, Γ(*s*), is the variance of the Poisson spikes, i.e., Γ(*s*) = diag(***r***(*s*)). We find (see Supplementary Information) that in this case the steady-state covariance Σ(*s*) is well approximated by the diagonal matrix 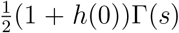 in other words, the covariance of the rates is dominated by the diagonal variance terms.

#### Decoding

The stochastic fluctuations in the neurons’ firing rates introduce a degree of noise that decreases the information content of each spike. To see this we look at the Fisher information of the neurons with fluctuating rates. Given a vector of rates 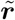, each Poisson neuron emits spikes independently (i.e., 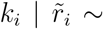 Poisson(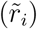)), but the rates themselves are stochastic, and sampled from a Gaussian distribution with covariance 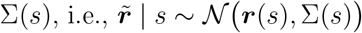. We approximate the distribution of the spike counts ***k*** conditional on a stimulus *s* with a matching-moments Gaussian distribution, i.e.,

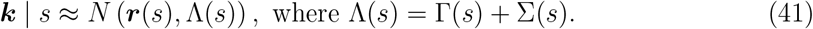

With the diagonal approximation of Σ mentioned above, we have Λ(*s*) ≈ *β* diag(***r***(*s*)), where 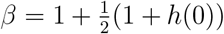. Note that 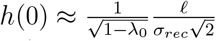. Thus if the connectivity strength is not too large (i.e., *λ*_0_ not too close to 1), and the network is relatively ‘dense’ (i.e., *ℓ* ≪*σ*_*rec*_) such that a neuron averages random inputs from many neighboring neurons — two conditions that our choice of parameters verifies — then *h*(0) is small, and thus *β* ≈ 3*/*2.

Hence the approximate linear Fisher information for neuron *i* is a fraction 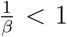 of its Fisher information in the absence of fluctuations, i.e.,

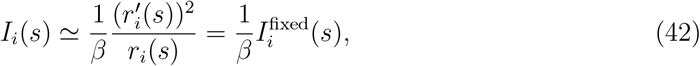

and the overall Fisher information is 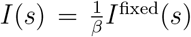. In other words, the network with fluctuating rates is equivalent to the network with fixed rates, but with the rates divided by *β*. Thus we choose the loss function

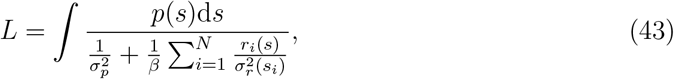

and the decoder we use in our simulations is the fixed-rate Bayesian-mean decoder (Eq. 10) with the spike counts *k*_*i*_ divided by *β*, i.e.,

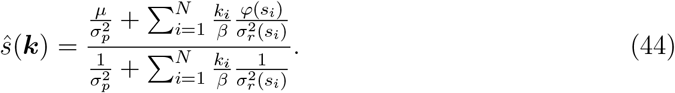

### Solving the efficient-coding problem

Using the continuous approximations derived above, we have

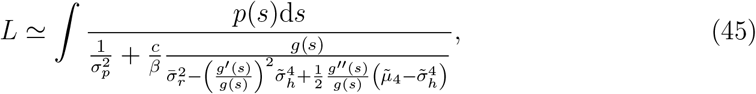

and

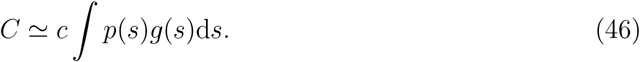

The Euler-Lagrange equation implied by Eq. 16 and *L* and *C* is not amenable to an analytic solution. Thus we adopt a crude approximation strategy, whereby wherever *g*(*s*) is nonzero, we consider that in the denominator of the integrand of *L* the precision term that comes from the prior, 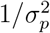, is small in comparison to the other term (which is the precision term that comes from the likelihood). Under this assumption the problem simplifies (see Supplementary Information) to minimizing the integral of the Lagrangian

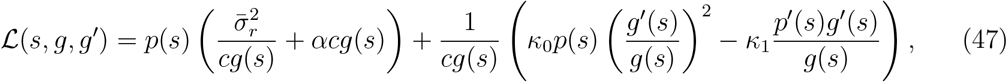

where *κ*_0_ and *κ*_1_ are constants. The two terms in the first parenthesis are readily interpreted. First, 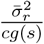 is the mean squared-error when the gains are uniform (*g*′ = 0): it scales as the squared-width of the tuning curves, and it is inversely related to the gains. Second, *αcg*(*s*) is the spiking cost, which increases proportionally to the gains. The second parenthesis captures the impact of the variations of the gains on the loss function. The first term increases with (*g*′)^2^ and is thus minimized by ‘flat’ gain profiles that spread out evenly the encoding resource. The second term, crucially, decreases with *p*′*g*′, and thus it favors gain profiles that allocate stronger gains towards the more frequent stimuli.

We find an approximate solution by positing small fluctuations of *g*(*s*), i.e., we assume that 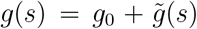, where 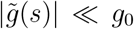. The expansion of *g*(*s*) in ℒ using this small-fluctuation assumption yields a simpler functional-minimization problem; in turn, the corresponding Euler-Lagrange equation 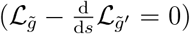 yields a differential equation for 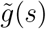 whose solution is dominated by a linear function of log *p*(*s*), resulting in our solution for the gain profile (Eq. 18). Note that with the Gaussian prior that we have assumed, the optimal gains take a quadratic form (bounded below by 0), as

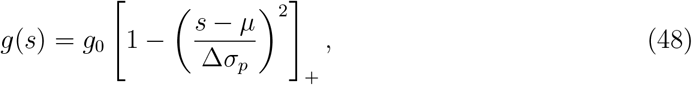

where *g*_0_ is the maximal gain in the network, while Δ determines the neurons whose gains are non-vanishing, as those whose feedforward locations *s* belongs in the interval [*µ* − *σ*_*p*_Δ, *µ* + *σ*_*p*_Δ]. The details of these derivations are given in Supplementary Information.

### Network optimization and simulations

To optimize the network’s gains we use the Adam optimizer as implemented in PyTorch 2.9.1. For each prior, we ran the same gradient-based optimization, starting from three distinct initializations: (i) the analytically approximate optimal gain profile smoothed with a kernel of width equal to the ‘typical’ tuning curve width (Eq. 33); (ii) a gain profile constant across the whole stimulus space, and equal to the mean of the analytic solution; and (iii) the same constant profile modulated by the prior (scaled to preserve the mean gain). For each initialization, optimization used Adam with an initial learning rate of 0.01, and a learning-rate ‘schedule’ (ReduceLROnPlateau) that halved the learning rate after 50 steps without sufficient improvement in the loss, with improvement defined by a relative threshold *ε* = 10^−6^. Optimization was terminated by early stopping after a fixed patience (500 steps without sufficient improvement) or upon reaching a maximum of 50,000 steps. The final gain profile was selected as the solution achieving the lowest attained loss across the three runs.

We ran simulations of the recurrently-connected Poisson neurons, to verify our calculations and to obtain the response variance shown in Fig. 5F. We set the network time constant to *τ* = 0.01 units of time, and for each value of the stimulus *s* we simulate the network 1000 times, for a duration *T* = 1, after a ‘burn-in’ period of duration 0.5, and with simulation time steps *dt* = 0.001. We start with rates equal to the gain-modulated feedforward tuning curves (***r***(*s, t* = 0) = *g* ⊙ *f* (*s*)). At each time step we sample random counts ***k*** from Poisson distributions with rates *r*_*i*_*dt*, and we then update the rates according to the dynamics equation (Eq. 1), where we set ***ξ***_*t*_*dt* = ***k*** − ***r****dt* so as to match the moments of the white noise ***ξ***_*t*_ while maintaining the Poisson stochasticity of the spikes. A comparison of the obtained empirical rates to the analytical rates (Eq. 2) is shown in Supplementary Information.

### Task and subjects

The experiment was a variant of the one presented in Ref. [10]. In the original design, each prior was tested in a separate block of trials, with the distribution remaining constant throughout each block. By contrast, here the prior changed randomly from trial to trial, enabling us to examine whether relatively quick changes in the stimulus distribution resulted in changes in subjects’ estimates. The experiment was pre-registered on AsPredicted (https://aspredicted.org/48yv-zs3c.pdf). The task was divided in two sections. In the first section, the three ranges were presented “separately”, i.e., in this section there were three blocks of successive trials, and each block featured a prior that did not change across trials. Subjects were explicitly informed of the prior, at the beginning of each block. In the second section, at each trial the prior was chosen randomly and explicitly told to the subject. Upon presentation of the prior cue, subjects could proceed in a self-paced fashion after a minimum of one second. The prior cue was then removed from the screen; then a fixation cross was shown for 500ms, after which the cloud of dots was shown for 500ms. In all trials the size of the slider stayed the same, and ranged from 25 to 95. Upon appearing on screen, the slider did not show any ‘thumb’, i.e., it had no preset initial value. Subjects could click anywhere on this empty slider; a thumb would then appear at this location, and the corresponding selected number was then shown. Subjects were then free to adjust their selected response before submitting.

Each section comprised three kinds of phases. In the ‘Learning’ phases (15 trials), the correct answer (that is, the correct number of dots) was directly shown to the subject, together with the cloud of dots. The only action required in this phase was to press the spacebar to move on to the next trial. In the ‘Feedback’ phases (30 trials), subjects were asked to estimate the number of dots in each cloud, after which feedback was provided, i.e., the correct number was shown. In the ‘No-Feedback’ phases (10 trials in each block of the first section, 150 trials in the second section), subjects estimated the number of dots, and no feedback was provided. The across-subject average number of trials with a Narrow prior was 50.6 (sd: 5.3), with a Medium prior, 49.5 (sd: 6.0), and with a Wide prior, 49.9 (sd: 5.7). Each dot had a radius of 10 pixels. The location of each dot was sampled from a bivariate Gaussian distribution with a standard deviation of 64 pixels (in both directions), conditional on dots not overlapping, and truncated to a display square with sides of 448 pixels.

The performance of subjects in both the Feedback phases and the No-Feedback phases impacted their reward. Data analysis was conducted only with the data from the No-Feedback phase of the second section. The reward comprised a $3 (USD) base pay and a performance bonus, calculated as follows. Subjects accumulated points throughout the experiment. The amount of points obtained in a trial was equal to 100(1 − ((ŝ − *s*)*/*12)^2^), where *s* was the correct number of dots and ŝ was the subject’s estimate. Every 1000 points obtained in the experiment was worth 40¢ in the performance bonus. Each trial in the Feedback and in the No-Feedback phases resulted in points. The points formula just presented allowed for negative numbers. In this case, it was deducted from the cumulated number of points. If at the end of the experiment a subject had a negative total number of points, then the performance bonus was set to zero, and the responses of this subject were not included in the analysis (as described in the pre-registration). This was the case for one subject.

A total of 60 subjects participated in the experiments (36 male, 24 female, average age: 43.1, standard deviation (sd): 11.8). The average performance bonus was $7.09 (sd: 1.89). The average duration of the experiment was 52 minutes (sd: 19.3 minutes). Subjects were recruited on Amazon Mechanical Turk through CloudResearch [65]. The study protocol was approved by the Institutional Review Board (IRB) of Harvard University (protocol IRB15-2048). The task was coded with jsPsych [66].

### Data analysis

The variance estimates presented in Figure 1B correspond to the posterior-mean estimates of the fixed-effect components of a statistical model that included subject-specific random effects. In other words, they are Bayesian estimates of the average variance across subjects. More precisely, the statistical model was specified by the following equations, where *x* is a number of dots, *c* is the condition (Narrow, Medium, or Wide), 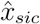 is the response of subject *s* in trial *i* of condition *c*, and *N*_+_ is the Gaussian distribution truncated to the positive numbers:

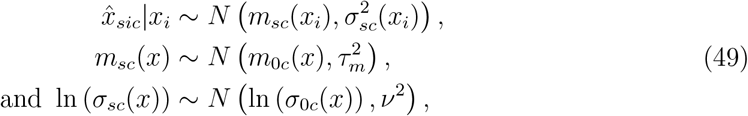

with the priors

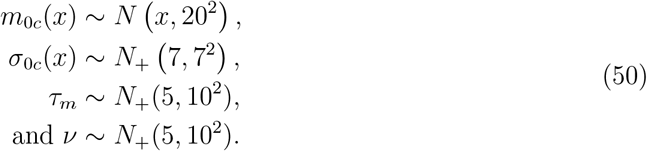

In other words, for a given number *x* presented in condition *c, m*_0*c*_(*x*) and *σ*_0*c*_(*x*) are the fixed-effect average and standard deviation of responses (while *m*_*sc*_(*x*) and *σ*_*sc*_(*x*) are the corresponding subject-specific random effects). This statistical model was estimated using Stan with the HMC-NUTS sampler [67] (10 chains of 1000 samples each, following 1000 warmup iterations.) We then compute (for each sample) the average across numbers of the response variance (in our analyses we also considered a model in which the response standard deviations, *σ*_0*c*_ and *σ*_*sc*_, do not depend on *x*; this model yields very similar estimates and prompts the same conclusions). The error bars in Figure 1B correspond to the 5th and 95th percentile of the posterior. The posterior probabilities that the variance is larger in the Narrow condition than in the Medium condition, and that it is larger in the Medium condition than in the Wide condition, are both lower than 10^−5^ (no sample supports these hypotheses). The fifth percentiles of the difference of standard deviations are 3.8 (Medium vs. Narrow) and 3.5 (Wide vs. Medium), i.e., with posterior probability 95% the differences are greater than these values. All other data analyses were conducted using NumPy and Scipy, and figures were made using Matplotlib [68–70].

## Data and code availability statements

The data for the behavioral experiment is publicly available at https://osf.io/vhdqe/. The code used for the model and the figures is publicly available at https://osf.io/wa7hm/.

## Acknowledgements

We thank Lyndon Duong for helpful discussions. This research was supported by the Kempner Institute for the Study of Natural and Artificial Intelligence and a Polymath Award from Schmidt Sciences (S.J.G.).

## Notes

https://osf.io/vhdqe/

https://osf.io/wa7hm/

### Competing Interest Statement

The authors have declared no competing interest.

### Summary of Updates

Model now includes noise. Connectivity matrix with global inhibition was examined. Bayesian model of behavior was implemented.

