## Supplementary Information for "Fast efficient coding and sensory adaptation in gain-adaptive recurrent networks"

Arthur Prat-Carrabin, Maximilian V. Harl, Samuel J. Gershman

February 8, 2026

#### Effective tuning curves: mathematical derivations

##### Eigenelements of $W$

We first diagonalize the matrix $W$ using the discrete cosine transform. Consider the vector $\boldsymbol{z}^{(0)} = (1, \ldots, 1)$. The $i^{th}$ element of the product $W\boldsymbol{z}^{(0)}$ is

$$\begin{aligned}
\sum_{j=1}^{N} W_{ij} z_j &= \sum_{j=1}^{N} W_{ij} \\
&= \lambda_0 \sum_{j=1}^{N} \frac{\ell}{\sigma_{rec}\sqrt{2\pi}} \exp\left(-\frac{(i-j)^2\ell^2}{2\sigma_{rec}^2}\right) \\
&\simeq \lambda_0 \int \frac{1}{\sigma_{rec}\sqrt{2\pi}} \exp\left(-\frac{(s_i - s_j)^2}{2\sigma_{rec}^2}\right) \mathrm{d}s_j \\
&\simeq \lambda_0,
\end{aligned} \tag{S1}$$

where we have approximated the discrete sum with a continuous integral. We thus obtain $W\boldsymbol{z}^{(0)} \simeq \lambda_0 \boldsymbol{z}^{(0)}$, i.e., $\boldsymbol{z}^{(0)}$ is (approximately) an eigenvector of $W$, with associated eigenvalue $\lambda_0$.

Similarly, the vectors $\boldsymbol{z}^{(k)} = \left(\cos\left(\frac{\pi k}{N}(i + \frac{1}{2})\right)\right)_{i=0,\ldots,N-1}$ are eigenvectors, with associated approximate eigenvalues $\lambda_k < \lambda_0$ defined as

$$\lambda_k = \lambda_0 \exp\left(-\frac{1}{2}\left(\pi \frac{\sigma_{rec} k}{\ell N}\right)^2\right), \tag{S2}$$

where $k = 0, \ldots, N-1$. Below we compare the eigenvalues of $W$ with their approximations (Fig. S6).

#### Approximations of $M$

We can use the diagonalization of $W$ to compute an approximation of $M = (I - W)^{-1}$. Specifically, $W = QDQ^T$, where $D = \mathrm{Diag}(\lambda_k)$ and $Q$ is an orthogonal matrix whose columns are the normalized eigenvectors of $W$. As for $M$, we thus have $M = Q(I - D)^{-1}Q^T$, where $(I - D)^{-1} = \mathrm{Diag}(\frac{1}{1-\lambda_k})$. Here we note a result that will be useful later: as $\boldsymbol{z}^{(0)}$ is an approximate eigenvector of $W$ with eigenvalue $\lambda_0$, we have $(I - W)\boldsymbol{z}^{(0)} = (1 - \lambda_0)\boldsymbol{z}^{(0)}$, and thus $\frac{1}{1-\lambda_0}\boldsymbol{z}^{(0)} = M\boldsymbol{z}^{(0)}$ (i.e., $\boldsymbol{z}^{(0)}$ is the eigenvector of $M$ associated with the eigenvalue $\frac{1}{1-\lambda_0}$); thus

$$\forall i,\ \sum_{j=1}^{N} M_{ij} = \frac{1}{1-\lambda_0}. \tag{S3}$$

As for the elements of $M$, we have $M = Q(I - D)^{-1}Q^T = \sum_{k=1}^{N} \frac{1}{1-\lambda_k} Q_{ik}Q_{jk}$, where $Q_{ik} = 1/\sqrt{N}$ if $k = 0$, otherwise $Q_{ik} = \sqrt{2/N}\cos\left(\frac{\pi k}{N}(i + \frac{1}{2})\right)$; and $\sum_{i=1}^{N} Q_{ik}Q_{il} = \delta_{kl}$. Thus

$$M_{ij} = \frac{1}{1-\lambda_0}\frac{1}{N} + \frac{2}{N}\sum_{k=1}^{N-1} \frac{1}{1-\lambda_k}\cos\left(\frac{\pi k}{N}(i+\frac{1}{2})\right)\cos\left(\frac{\pi k}{N}(j+\frac{1}{2})\right). \tag{S4}$$

Assuming that $N$ is large and relying again on a continuous approximation of this discrete sum, we find that $M_{ij}$ can be approximated by an infinite sum of Gaussian functions, as

$$M_{ij} \simeq \delta_{ij} + \ell \sum_{m=1}^{\infty} \frac{\lambda_0^m}{\sigma_{rec}\sqrt{m}\sqrt{2\pi}} \exp\left(-\frac{(i-j)^2\ell^2}{2m\sigma_{rec}^2}\right), \tag{S5}$$

where $\delta_{ij}$ is the Kronecker delta. We have used the equalities $\frac{1}{1-\lambda_k} = \sum_{m=0}^{\infty} \lambda_k^m$ and $\int_0^\infty e^{-ax^2/2}\cos(bx)\mathrm{d}x = \sqrt{\frac{\pi}{2a}}e^{-\frac{b^2}{2a}}$.

If $\lambda_0$ is small (e.g. $\lambda = 0.15$), then a good approximation of $M$ can be obtained by keeping just the first Gaussian function in the sum, i.e.,

$$M_{ij} \simeq \delta_{ij} + \lambda_0 \frac{\ell}{\sigma_{rec}\sqrt{2\pi}} \exp\left(-\frac{(i-j)^2\ell^2}{2\sigma_{rec}^2}\right). \tag{S6}$$

For larger values of $\lambda_0$, more Gaussian terms are necessary to obtain a reasonably good approximation. We call this approximation (with only one Gaussian function) the 'Gaussian approximation'. Note that this approximation is $M_{ij} \simeq \delta_{ij} + W_{ij}$. Hence it is equivalent to noting that $(I - W)^{-1} = \sum_{m=0}^{\infty} W^m$, and keeping only the first two terms of this series.

We can also further approximate Eq. S5 with yet another continuous integral, and use the equality $\int_0^\infty \frac{1}{\sqrt{m}}\exp(-\alpha m - \beta/m)\mathrm{d}m = \sqrt{\frac{\pi}{\alpha}}e^{-2\sqrt{\alpha\beta}}$. We obtain

$$M_{ij} = \delta_{ij} + \frac{1}{\ln(1/\lambda_0)}\frac{\ell}{2\nu_{rec}} \exp\left(-\frac{|i-j|\ell}{\nu_{rec}}\right), \tag{S7}$$

where

$$\nu_{rec} = \frac{\sigma_{rec}}{\sqrt{2\ln(1/\lambda_0)}}. \tag{S8}$$

We call this approximation the 'Laplace approximation'. We find that it is reasonably good if $\sigma_{rec}$ is small (e.g., $\sigma_{rec} = \ell N/100$). For larger $\sigma_{rec}$ (e.g., $\sigma_{rec} = \ell N/20$), the approximation is reasonable if $\lambda_0$ is large (e.g., $\lambda_0 = 0.85$). For small $\lambda_0$, it is better to use the Gaussian approximation defined above. Below we provide comparisons of the elements of $M - I$ with these various approximations (Figs. S7-S8).

In short, roughly, the Gaussian approximation is good for small $\lambda_0$, and the Laplace approximation is preferable for larger $\lambda_0$. We find that interpolating between these two approximations, with a weight $\lambda_0$ on the Laplace approximation and a weight $1 - \lambda_0$ on the Gaussian approximation, provides a reasonably accurate solution for the whole range of $\lambda_0$. Specifically,

$$M_{ij} \simeq \delta_{ij} + h(s_i - s_j), \tag{S9}$$

where $h$ is the unimodal symmetric function

$$h(z) = (1-\lambda_0)\lambda_0 \frac{\ell}{\sigma_{rec}\sqrt{2\pi}} \exp\left(-\frac{z^2}{2\sigma_{rec}^2}\right) + \frac{\lambda_0}{\ln(1/\lambda_0)} \frac{\ell}{2\nu_{rec}} \exp\left(-\frac{|z|}{\nu_{rec}}\right). \tag{S10}$$

In what follows it will also be useful to consider the quantity

$$\bar{h} \equiv \frac{1}{\ell}\int h(z)\mathrm{d}z = (1-\lambda_0)\lambda_0 + \frac{\lambda_0}{\ln(1/\lambda_0)}, \tag{S11}$$

the squared-width quantity

$$\begin{aligned} \sigma_h^2 \equiv \frac{1}{\ell}\int z^2 h(z)\mathrm{d}z &= (1-\lambda_0)\lambda_0\sigma_{rec}^2 + \frac{\lambda_0}{\ln(1/\lambda_0)}\nu_{rec}^2 \\ &= \sigma_{rec}^2\lambda_0\left((1-\lambda_0) + \frac{1}{2\ln(1/\lambda_0)^2}\right), \end{aligned} \tag{S12}$$

and the 'fourth-moment' quantity

$$\mu_4 \equiv \frac{1}{\ell}\int z^4 h(z)\mathrm{d}z = (1-\lambda_0)\lambda_0 3\sigma_{rec}^4 + \frac{\lambda_0}{\ln(1/\lambda_0)} 24\nu_{rec}^4 \tag{S13}$$

$$= 3\sigma_{rec}^4\lambda_0\left((1-\lambda_0) + \frac{2}{(\ln(1/\lambda_0))^3}\right), \tag{S14}$$

as well as their values normalized by $1 + \bar{h}$,

$$\tilde{\sigma}_h^2 \equiv \frac{\sigma_h^2}{1+\bar{h}} \text{ and } \tilde{\mu}_4 \equiv \frac{\mu_4}{1+\bar{h}}. \tag{S15}$$

### Stationary noise covariance

The stationary covariance $\Sigma$ of the linear stochastic dynamics satisfies the continuous Lyapunov equation

$$A\Sigma + \Sigma A + \Gamma = 0, \tag{S16}$$

where $A = W - I = -M^{-1}$. The solution is known in the integral representation

$$\Sigma = \int_0^\infty e^{tA}\,\Gamma\,e^{tA}\,dt. \tag{S17}$$

With the diagonal matrix $\Gamma = \mathrm{diag}(\boldsymbol{r})$, the entries of $\Sigma$ are

$$\Sigma_{mn} = \sum_k r_k \int_0^\infty [e^{tA}]_{mk}\,[e^{tA}]_{nk}\,dt. \tag{S18}$$

Note that $e^{tA} = e^{-t}\sum_{i\geq 0}\frac{t^i}{i!}W^i$ (the matrix element $[e^{tA}]_{mk}$ can be interpreted as the activity at neuron $m$ generated by neuron $k$ after a Poisson-distributed number of recurrent steps with mean $t$). Thus $[e^{tA}]_{mk}$ is appreciable only when $k$ lies within a neighborhood of $m$, and the product $[e^{tA}]_{mk}[e^{tA}]_{nk}$ is significant only in the region where the neighborhoods of $m$ and $n$ overlap. The sum over $k$ therefore acts as a local average of $r_k$ over this overlap region. Assuming slow variations of $r_k$, we use the approximation

$$r_k \approx \frac{r_m + r_n}{2}, \tag{S19}$$

which gives

$$\Sigma_{mn} \approx \frac{r_m + r_n}{2}\int_0^\infty \sum_k [e^{tA}]_{mk}[e^{tA}]_{nk}\mathrm{d}t \tag{S20}$$

$$= \frac{r_m + r_n}{2}\int_0^\infty [e^{tA}e^{tA}]_{mn}\mathrm{d}t = \frac{r_m + r_n}{2}\int_0^\infty [e^{2tA}]_{mn}\,dt \tag{S21}$$

$$= \frac{r_m + r_n}{2}\Big[-\frac{1}{2}A^{-1}\Big]_{mn} \tag{S22}$$

$$= \frac{r_m + r_n}{4}M_{mn}. \tag{S23}$$

Thus the stationary covariance matrix can be approximated as

$$\Sigma \approx \frac{1}{4}(M\Gamma + \Gamma M). \tag{S24}$$

The matrix $\Gamma$ is diagonal and $M$ is dominated by its diagonal elements $M_{nn} = 1 + h(0)$ (see $M - I$ in Figs. S7-S8), thus we further approximate $\Sigma$ as

$$\Sigma \approx \frac{1}{2}(1 + h(0))\Gamma. \tag{S25}$$

We find that this is a reasonable approximation with the chosen parameters of the network. Figure S9, below, compares the solution of the Lyapunov equation to the approximations in Eqs. S24 and S25.

#### Continuous approximations

Because we consider the case of relatively wide priors, we assume that the various functions characterizing the network ($g(s)$, $\varphi(s)$, etc.) vary slowly on the scale of the neurons' widths. We obtain the following useful continuous approximations. First, the effective location of neuron $i$ is

$$\varphi(s_i) \simeq \sum_j \gamma_{ij} s_j = s_i + \frac{\sum_j g_j h(s_i - s_j)(s_j - s_i)}{g_i + \sum_j g_j h(s_i - s_j)}. \tag{S26}$$

In the limit of a continuum of neurons,

$$\varphi(s) \simeq s + \frac{\int g(s_j)\, h(s - s_j)(s_j - s) \frac{\mathrm{d}s_j}{\ell}}{g(s) + \int g(s_j) h(s - s_j) \frac{\mathrm{d}s_j}{\ell}} \simeq s + \frac{g'(s)\sigma_h^2}{g(s)(1+\bar{h})} \tag{S27}$$

$$= s + \frac{g'(s)}{g(s)} \tilde{\sigma}_h^2, \tag{S28}$$

which is Eq. 17. Hence, with uniform gains ($g' = 0$), $\varphi$ reduces to the identity function ($\varphi(s) = s$). The effective squared-width of neuron $i$ is

$$\sigma_r^2(s_i) = \sigma_f^2 + \sum_j \gamma_{ij} (s_j - \varphi(s_i))^2 \tag{S29}$$

$$= \sigma_f^2 - (\varphi(s_i) - s_i)^2 + \sum_j \gamma_{ij} (s_j - s_i)^2, \tag{S30}$$

where

$$\sum_j \gamma_{ij} (s_j - s_i)^2 = \frac{\sum_j g_j h(s_i - s_j)(s_j - s_i)^2}{g_i + \sum_j g_j h(s_i - s_j)}. \tag{S31}$$

A continuous approximation of this quantity is

$$\frac{\int g(s_j) h(s - s_j)(s - s_j)^2 \frac{\mathrm{d}s_j}{\ell}}{g(s) + \int g(s_j) h(s - s_j) \frac{\mathrm{d}s_j}{\ell}} \simeq \frac{g(s)\sigma_h^2 + \frac{1}{2} g''(s)\mu_4 + O(\partial^4 g)}{g(s) + g(s)\bar{h} + \frac{1}{2} g''(s)\sigma_h^2 + O(\partial^4 g)} \tag{S32}$$

$$\simeq \frac{\tilde{\sigma}_h^2 + \frac{1}{2}\frac{g''(s)}{g(s)}\frac{\mu_4}{1+\bar{h}} + O(\partial^4 g/g)}{1 + \frac{1}{2}\frac{g''(s)}{g(s)}\tilde{\sigma}_h^2 + O(\partial^4 g/g)} \tag{S33}$$

$$\simeq \left(\tilde{\sigma}_h^2 + \frac{1}{2}\frac{g''(s)}{g(s)}\tilde{\mu}_4 + O(\partial^4 g/g)\right)\left(1 - \frac{1}{2}\frac{g''(s)}{g(s)}\tilde{\sigma}_h^2 + O(\partial^4 g/g)\right) \tag{S34}$$

$$\simeq \tilde{\sigma}_h^2 + \frac{1}{2}\frac{g''(s)}{g(s)}\left(\tilde{\mu}_4 - \tilde{\sigma}_h^4\right). \tag{S35}$$

Thus

$$\sigma_r^2(s) = \bar{\sigma}_r^2 - \left(\frac{g'(s)}{g(s)}\right)^2 \tilde{\sigma}_h^4 + \frac{1}{2}\frac{g''(s)}{g(s)}\left(\tilde{\mu}_4 - \tilde{\sigma}_h^4\right), \tag{S36}$$

where we denote by $\bar{\sigma}_r^2$ the typical width of the tuning curves:

$$\bar{\sigma}_r^2 = \sigma_f^2 + \tilde{\sigma}_h^2. \tag{S37}$$

The total expected spike count, given a stimulus $s$, is

$$\sum_{i=1}^{N} r_i(s) = \sum_{i=1}^{N}\sum_{j=1}^{N} M_{ij} g_j f_j(s) = \sum_{j=1}^{N} g_j f_j(s) \sum_{i=1}^{N} M_{ij} \tag{S38}$$

$$= \frac{1}{1-\lambda_0}\sum_{i=1}^{N} g_j f_j(s) \simeq \frac{1}{1-\lambda_0}\int g(s_j)\exp\left(-\frac{(s-s_j)^2}{2\sigma_f^2}\right)\frac{\mathrm{d}s_j}{\ell} \tag{S39}$$

$$\simeq \frac{\sigma_f\sqrt{2\pi}}{\ell(1-\lambda_0)} g(s) \tag{S40}$$

$$= cg(s), \tag{S41}$$

where we have used Eq. S3, and where

$$c = \frac{\sigma_f\sqrt{2\pi}}{\ell(1-\lambda_0)}. \tag{S42}$$

The expected precision of the decoder (Eq. 30) is driven by the quantity

$$\sum_{i=1}^{N}\frac{r_i(s)}{\sigma_r^2(s_i)} \simeq \int \frac{r_i(s)}{\sigma_r^2(s_i)}\frac{\mathrm{d}s_i}{\ell} \simeq \frac{1}{\sigma_r^2(\varphi^{-1}(s))}\int r_i(s)\frac{\mathrm{d}s_i}{\ell} \tag{S43}$$

$$\simeq \frac{1}{\sigma_r^2(\varphi^{-1}(s))}\sum_{i=1}^{N} r_i(s) \simeq c\frac{g(s)}{\sigma_r^2(\varphi^{-1}(s))} \tag{S44}$$

$$\simeq c\frac{g(s)}{\sigma_r^2(s_i)}. \tag{S45}$$

As for the Fisher information,

$$I(s) \simeq \frac{1}{\beta}\sum_i \frac{(r_i'(s))^2}{r_i(s)} \simeq \frac{1}{\beta}\sum_i r_i(s)\frac{(s-\varphi(s_i))^2}{\sigma_r^4(s_i)} \tag{S46}$$

$$\simeq \frac{1}{\beta}\frac{1}{\ell}\int G(s_i)\frac{\sigma_f}{\sigma_r^5(s_i)}\exp\left(-\frac{(s-\varphi(s_i))^2}{\sigma_r^2(s_i)}\right)(s-\varphi(s_i))^2\mathrm{d}s_i \tag{S47}$$

$$= \frac{1}{\beta}\frac{1}{\ell}\int G(\varphi^{-1}(z))\frac{\sigma_f}{\sigma_r^5(\varphi^{-1}(z))}\exp\left(-\frac{(s-z)^2}{\sigma_r^2(\varphi^{-1}(z))}\right)(s-z)^2(\varphi^{-1})'(z)\mathrm{d}z \tag{S48}$$

$$\simeq \frac{1}{\beta}\frac{1}{\ell}G(\varphi^{-1}(s))\frac{\sigma_f\sqrt{2\pi}}{\sigma_r^2(\varphi^{-1}(s))}(\varphi^{-1})'(s) \tag{S49}$$

$$\simeq \frac{c}{\beta}\frac{g(s)}{\sigma_r^2(s_i)}. \tag{S50}$$

### Solving the efficient-coding problem

We now consider the efficient-coding problem defined in Eq. 16. Using the continuous approximations just derived, we have

$$L \simeq \int \frac{p(s)\mathrm{d}s}{\frac{1}{\sigma_p^2} + \frac{c}{\beta}\frac{g(s)}{\sigma_r^2(s_i)}} \tag{S51}$$

$$\simeq \int \frac{p(s)\mathrm{d}s}{\frac{1}{\sigma_p^2} + \frac{c}{\beta}\frac{g(s)}{\bar{\sigma}_r^2 - \left(\frac{g'(s)}{g(s)}\right)^2\tilde{\sigma}_h^4 + \frac{1}{2}\frac{g''(s)}{g(s)}\left(\tilde{\mu}_4 - \tilde{\sigma}_h^4\right)}}, \tag{S52}$$

and

$$C \simeq c\int p(s)g(s)\mathrm{d}s. \tag{S53}$$

The Euler-Lagrange equation implied by $L$ and $C$ is not amenable to an analytic solution. Thus we adopt a crude approximation strategy, whereby wherever $g(s)$ is not zero, we consider that in the denominator of the integrand of $L$ the precision term that comes from the prior, $1/\sigma_p^2$, is small in comparison to the other term (which is the precision term that comes from the likelihood). Specifically, let $S$ be the set where $g$ is non-vanishing, i.e., $S = \{s : g(s) > 0\}$. On $S$, we neglect $1/\sigma_p^2$ in the loss function, and thus the loss on $S$ is

$$L_S = \beta\int_S p(s)\frac{\bar{\sigma}_r^2 - \left(\frac{g'(s)}{g(s)}\right)^2\tilde{\sigma}_h^4 + \frac{1}{2}\frac{g''(s)}{g(s)}\left(\tilde{\mu}_4 - \tilde{\sigma}_h^4\right)}{cg(s)}\mathrm{d}s \tag{S54}$$

$$= \frac{\beta}{c}\int_S p(s)\left(\frac{\bar{\sigma}_r^2}{g(s)} - \frac{(g'(s))^2}{g(s)^3}\tilde{\sigma}_h^4 + \frac{1}{2}\frac{g''(s)}{g(s)^2}\left(\tilde{\mu}_4 - \tilde{\sigma}_h^4\right)\right)\mathrm{d}s. \tag{S55}$$

Integrating by parts,

$$\int_S p(s)\frac{g''(s)}{g(s)^2}\mathrm{d}s \simeq -\int_S p'(s)\frac{g'(s)}{g(s)^2}\mathrm{d}s + 2\int_S p(s)\frac{(g'(s))^2}{g(s)^3}\mathrm{d}s, \tag{S56}$$

thus

$$L_S = \frac{\beta}{c}\int_S \left[p(s)\left(\frac{\bar{\sigma}_r^2}{g(s)} + \frac{(g'(s))^2}{g(s)^3}(\tilde{\mu}_4 - 2\tilde{\sigma}_h^4)\right) - \frac{1}{2}(\tilde{\mu}_4 - \tilde{\sigma}_h^4)p'(s)\frac{g'(s)}{g(s)^2}\right]\mathrm{d}s. \tag{S57}$$

We thus seek to minimize the integral over $S$ of the Lagrangian

$$\mathcal{L}(s, g, g') = p(s)\left(\frac{\beta\bar{\sigma}_r^2}{cg(s)} + \alpha cg(s)\right) + \frac{\beta}{cg(s)}\left(\kappa_0 p(s)\left(\frac{g'(s)}{g(s)}\right)^2 - \kappa_1\frac{p'(s)g'(s)}{g(s)}\right), \tag{S58}$$

where $\bar{\sigma}_r^2 = \sigma_f^2 + \tilde{\sigma}_h^2$ is the typical squared-width of the tuning curves, and $\kappa_0$ and $\kappa_1$ are constants.

We find an approximate solution by positing small fluctuations of $g(s)$, i.e., we assume that $g(s) = g_0 + \tilde{g}(s)$, where $|\tilde{g}(s)| \ll g_0$. The expansion of $g(s)$ in $\mathcal{L}$ using this small-fluctuation assumption yields the following optimization problem for $\tilde{g}(s)$:

$$\min_{\tilde{g}} \int_S \mathcal{L}_2(s, \tilde{g}, \tilde{g}')\mathrm{d}s, \tag{S59}$$

where

$$\mathcal{L}_2(s,\tilde{g},\tilde{g}') = p(s)\Big(\alpha c - \frac{\beta\bar{\sigma}_r^2}{cg_0^2}\Big)\tilde{g}(s) + \frac{\beta}{cg_0}\left((\tilde{\mu}_4 - 2\tilde{\sigma}_h^4)\frac{p(s)}{g_0^2}(\tilde{g}'(s))^2 - \frac{\tilde{\mu}_4 - \tilde{\sigma}_h^4}{2g_0}p'(s)\tilde{g}'(s)\right). \tag{S60}$$

Calculus of variations shows that solutions to such functional-minimization problem necessarily obey the Euler-Lagrange equation,

$$\frac{\partial\mathcal{L}_2}{\partial\tilde{g}} - \frac{\mathrm{d}}{\mathrm{d}s}\frac{\partial\mathcal{L}_2}{\partial\tilde{g}'} = 0. \tag{S61}$$

We have

$$\frac{\partial\mathcal{L}_2}{\partial\tilde{g}} = p(s)\Big(\alpha c - \frac{\beta\bar{\sigma}_r^2}{cg_0^2}\Big), \tag{S62}$$

and

$$\frac{\partial\mathcal{L}_2}{\partial\tilde{g}'} = 2(\tilde{\mu}_4 - 2\tilde{\sigma}_h^4)\frac{\beta p(s)}{cg_0^3}\tilde{g}'(s) - \beta\frac{\tilde{\mu}_4 - \tilde{\sigma}_h^4}{2cg_0^2}p'(s). \tag{S63}$$

We note that the Legendre condition is verified, that is,

$$\frac{\partial^2\mathcal{L}_2}{\partial\tilde{g}'^2} = 2(\tilde{\mu}_4 - 2\tilde{\sigma}_h^4)\frac{\beta p(s)}{cg_0^3} > 0, \tag{S64}$$

which implies that if a function is solution to the Euler-Lagrange equation, then it is a minimum of the functional. We now write the Euler-Lagrange equation, as

$$p(s)\Big(\alpha c - \frac{\beta\bar{\sigma}_r^2}{cg_0^2}\Big) = \frac{\mathrm{d}}{\mathrm{d}s}\left[2(\tilde{\mu}_4 - 2\tilde{\sigma}_h^4)\frac{\beta p(s)}{cg_0^3}\tilde{g}'(s) - \beta\frac{\tilde{\mu}_4 - \tilde{\sigma}_h^4}{2cg_0^2}p'(s)\right]. \tag{S65}$$

Integrating yields

$$C_0 + P(s)\Big(\alpha c - \frac{\beta\bar{\sigma}_r^2}{cg_0^2}\Big) = 2(\tilde{\mu}_4 - 2\tilde{\sigma}_h^4)\frac{\beta p(s)}{cg_0^3}\tilde{g}'(s) - \beta\frac{\tilde{\mu}_4 - \tilde{\sigma}_h^4}{2cg_0^2}p'(s), \tag{S66}$$

where $C_0$ is an integration constant. Given the symmetry of the prior we seek a solution $\tilde{g}$ that is even around $\mu$ (and $\tilde{g}'$, odd around $\mu$). This determines $C_0$. We then obtain an expression for the derivative of $\tilde{g}$, as

$$\tilde{g}'(s) = \frac{g_0}{2(\tilde{\mu}_4 - 2\tilde{\sigma}_h^4)\beta}\left[\beta\frac{\tilde{\mu}_4 - \tilde{\sigma}_h^4}{2}\frac{p'(s)}{p(s)} + \frac{\frac{1}{2} - P(s)}{p(s)}\Big(\beta\bar{\sigma}_r^2 - \alpha c^2 g_0^2\Big)\right]. \tag{S67}$$

Integrating, we obtain

$$\tilde{g}(s) = C_2 + \frac{g_0}{2(\tilde{\mu}_4 - 2\tilde{\sigma}_h^4)\beta}\left[\beta\frac{\tilde{\mu}_4 - \tilde{\sigma}_h^4}{2}\log\frac{p(s)}{p(\mu)} + \Big(\beta\bar{\sigma}_r^2 - \alpha c^2 g_0^2\Big)\int_\mu^s \frac{\frac{1}{2} - P(t)}{p(t)}\mathrm{d}s\right] \tag{S68}$$

We note that

$$\frac{\frac{1}{2} - P(t)}{p(t)} \approx \frac{\frac{1}{2} - P(\mu) - p(\mu)(t-\mu)}{p(\mu)} = \mu - t \tag{S69}$$

and thus the second term in the brackets is approximately quadratic in $s-\mu$, as

$$\int_{\mu}^{s} \frac{\frac{1}{2}-P(t)}{p(t)}\mathrm{d}s \approx -\frac{1}{2}(s-\mu)^2. \tag{S70}$$

As for the first term in the brackets, for $s$ close to $\mu$ we have

$$\log p(s) \approx \log p(\mu) + \frac{p''(\mu)}{2p(\mu)}(s-\mu)^2 \approx \log p(\mu) - \frac{(s-\mu)^2}{2\,\mathrm{Var}[p]}, \tag{S71}$$

with strict equality for a Gaussian distribution. Thus the sum of terms in the brackets is approximately Gaussian. Comparing the orders of magnitude of these two terms, we find that with our parameters, $\beta\frac{\tilde{\mu}_4-\tilde{\sigma}_h^4}{2}\frac{1}{\mathrm{Var}[p]}$ is significantly larger than $|\beta\bar{\sigma}_r^2-\alpha c^2 g_0^2|$. Hence we keep only the first term, and obtain $\tilde{g}(s)$ as a linear function of $\log p(s)$, resulting in our solution for the gain profile (Eq. 18).

With the Gaussian prior that we have assumed, the optimal gains take a quadratic form (bounded below by 0), as

$$g(s) = g_0\left[1-\left(\frac{s-\mu}{\Delta\sigma_p}\right)^2\right]_+, \tag{S72}$$

where $g_0>0$ and $\Delta>0$ are two parameters chosen to optimize Eq. S59. The parameter $g_0$ is the maximal gain in the network, while $\Delta$ determines the neurons whose gains are non-vanishing, as those whose feedforward locations $s$ belongs in the interval $[\mu-\sigma_p\Delta, \mu+\sigma_p\Delta]$. Finally, this solution implies that the variations of the sum of the rates, $\sum r_i(s)$ (directly related to the gains through Eq. 34) occur on a wide scale comparable to that of the prior, justifying our neglecting this term in deriving an approximation to the posterior.

### Fisher information of a continuum of neurons

We consider a population of $N$ Poisson neurons with Gaussian tuning curves with locations $s_i$, width $\sigma_i$, and gain $g_i$, i.e., the firing rate of neuron $i$ as a function of the stimulus is

$$\rho_i(s) = g_i \exp\left(-\frac{(s-s_i)^2}{2\sigma_i^2}\right). \tag{S73}$$

The Fisher information with respect to $s$ of neuron $i$ is

$$I_i(s) = \frac{(\rho_i'(s))^2}{\rho_i(s)} = \frac{(s-s_i)^2}{\sigma_i^4}g_i\exp\left(-\frac{(s-s_i)^2}{2\sigma_i^2}\right). \tag{S74}$$

The total Fisher information of the population, if neurons fire independently, is the sum $I(s)=\sum I_i(s)$, which we approximate with an integral, as

$$I(s) \simeq \int \frac{(s-\tilde{s})^2}{\sigma(\tilde{s})^4}g(\tilde{s})\exp\left(-\frac{(s-\tilde{s})^2}{2\sigma(\tilde{s})^2}\right)d(\tilde{s})\mathrm{d}\tilde{s}, \tag{S75}$$

where $d(\tilde{s})$ is the density and $\tilde{s}$, $\sigma(\tilde{s})$, and $g(\tilde{s})$ replace the discrete tuning curves' parameters $s_i$, $\sigma_i$, and $g_i$. Assuming that $g(\tilde{s})$, $\sigma(\tilde{s})$, and $d(\tilde{s})$ change slowly on the scale of $\sigma(\tilde{s})$, this is approximately

$$I(s) \simeq \sqrt{2\pi}\frac{g(s)d(s)}{\sigma(s)}. \tag{S76}$$

In the recurrent network we are studying, the number of neurons whose effective locations are between $s$ and $s + \mathrm{d}s$ is $(\varphi^{-1}(s + \mathrm{d}s) - \varphi^{-1}(s))\frac{1}{\ell} = (\varphi^{-1})'(s)\frac{\mathrm{d}s}{\ell}$, and thus the density of neurons is $d(s) = (\varphi^{-1})'(s)\frac{1}{\ell}$.

### Inhibition

We consider a variant of our model in which global inhibition is added to the connectivity. Specifically, we add to the recurrence matrix $W$ a uniform lateral-inhibition term modeled by the rank-1 matrix $W^I$, where

$$W^I_{ij} = -\frac{J_I}{N}, \tag{S77}$$

where $J_I > 0$ is the inhibition strength. We set this parameter to 0.1, representing a background of moderate inhibition (compared to the recurrence strength, $\lambda_0 = 0.95$). All the other parameters of the network are kept identical. We optimize the network's gains to the Narrow, Medium, and Wide priors, and to the Control and Adaptation priors. Comparing the network's tuning curves with and without inhibition, we find that the inhibition suppresses the long tails of the curves, resulting in more realistic curve shapes (Fig. S1). In addition, we find that the network with inhibition also yields prior attraction: in the Narrow condition the neurons' locations align well with the prediction of efficient coding, while the prior attraction is less strong in the Medium condition (Fig. S2; compare to Fig. 4). Finally, we observe all the patterns found in the adaptation paradigm, including the repulsion and widening of the curves near the adapter, and the narrowing and attraction further from it (Fig. S3; compare to Fig. 6).

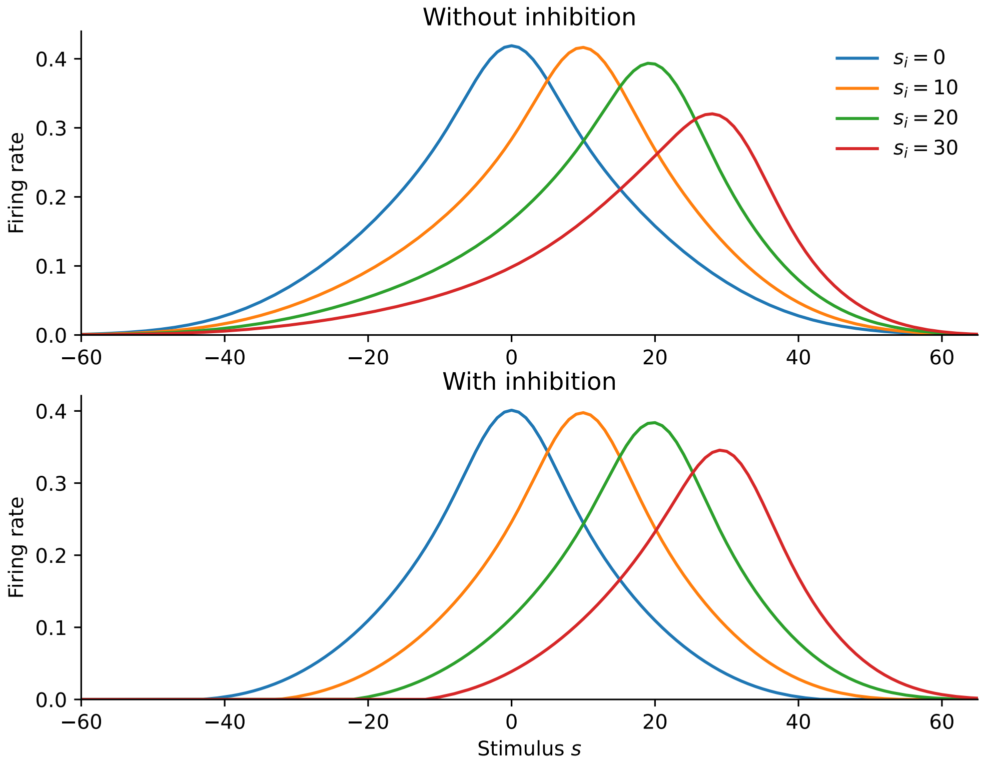


**Fig. S1: Tuning curves with and without inhibition.** Tuning curves of the neurons with feedforward locations 0, 10, 20, and 30, without inhibition (top) and with inhibition (bottom), with in each case the network's gains optimized to the Medium prior ($\sigma_p$)

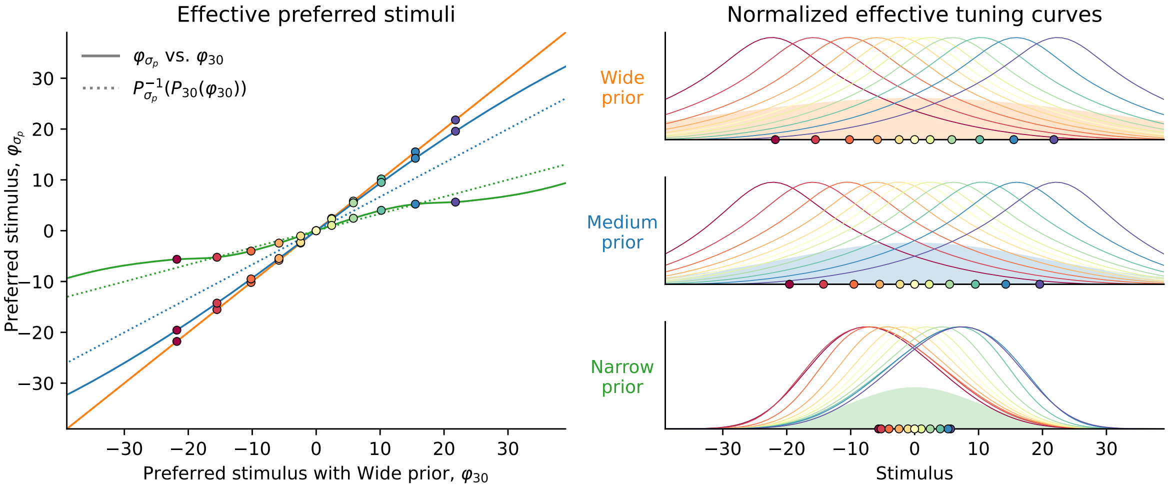


**Fig. S2: Prior attraction with network featuring global inhibition.** Compare with Fig. 4C,D.

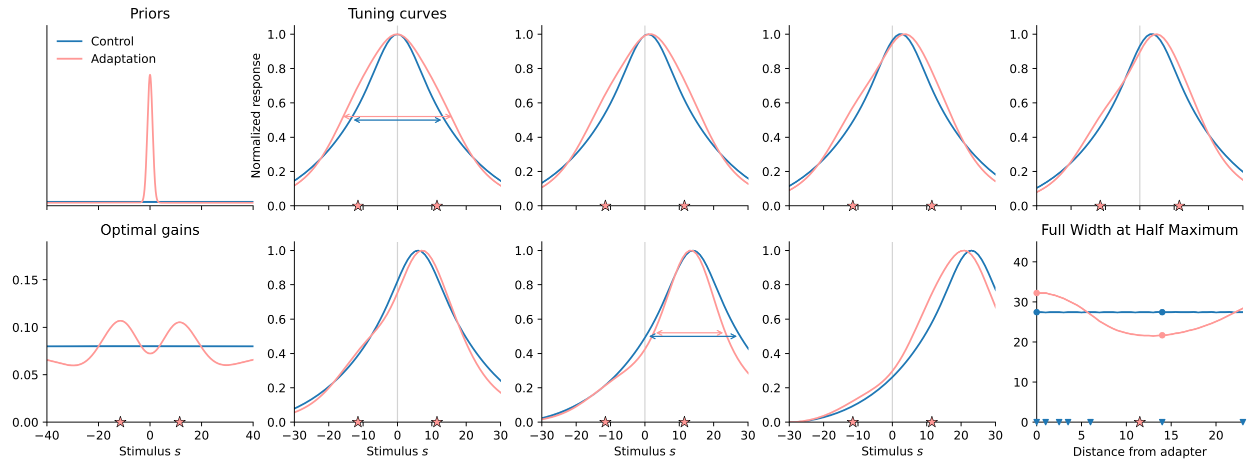


**Fig. S3: Adapter repulsion with network featuring global inhibition.** Compare with Fig. 6.

### Prior sweep

The adaptation prior in the main text is a mixture of 80% of the control prior (a uniform prior) and 20% of a Gaussian distribution with width $\sigma_p = 1$. Here we vary these parameters and optimize the network's gains to each resulting prior. We consider mixtures of 80/20, 40/60, and 10/90, with $\sigma_p = 1$, 5, and 10 (note that 0/100 with $\sigma_p = 10$ would correspond to the Narrow prior). Figure S4 shows the resulting optimal gains. The top left panel corresponds to the adaptation prior we have examined in the main text. When the Gaussian component of the prior narrows (going from right to left in the Figure), the bimodal 'M shape' of the gain profile accentuates. Conversely when the proportion of the uniform distribution in the prior increases (going from bottom to top), the gain profile flattens. We note in addition that a bimodality can already be seen with the relatively wide prior with $\sigma_p = 10$. Overall, we conclude that the gain profile we obtain with the adaptation prior, with maxima on the flanks of the prior peak, is a result robust to changes in the width and in the prominence of the peak in the prior.

### Uncertainty

Figure S5 shows the approximate standard deviation of the decoding posterior as a function of the prior width. Specifically, the posterior precision is approximately

$$\hat{\tau}(\boldsymbol{k}) = \frac{1}{\sigma_p^2} + \sum_{i=1}^{N} \frac{k_i}{\beta} \frac{1}{\sigma_r^2(s_i)}, \tag{S78}$$

and Figure S5 shows the average of the inverse square root of this quantity.

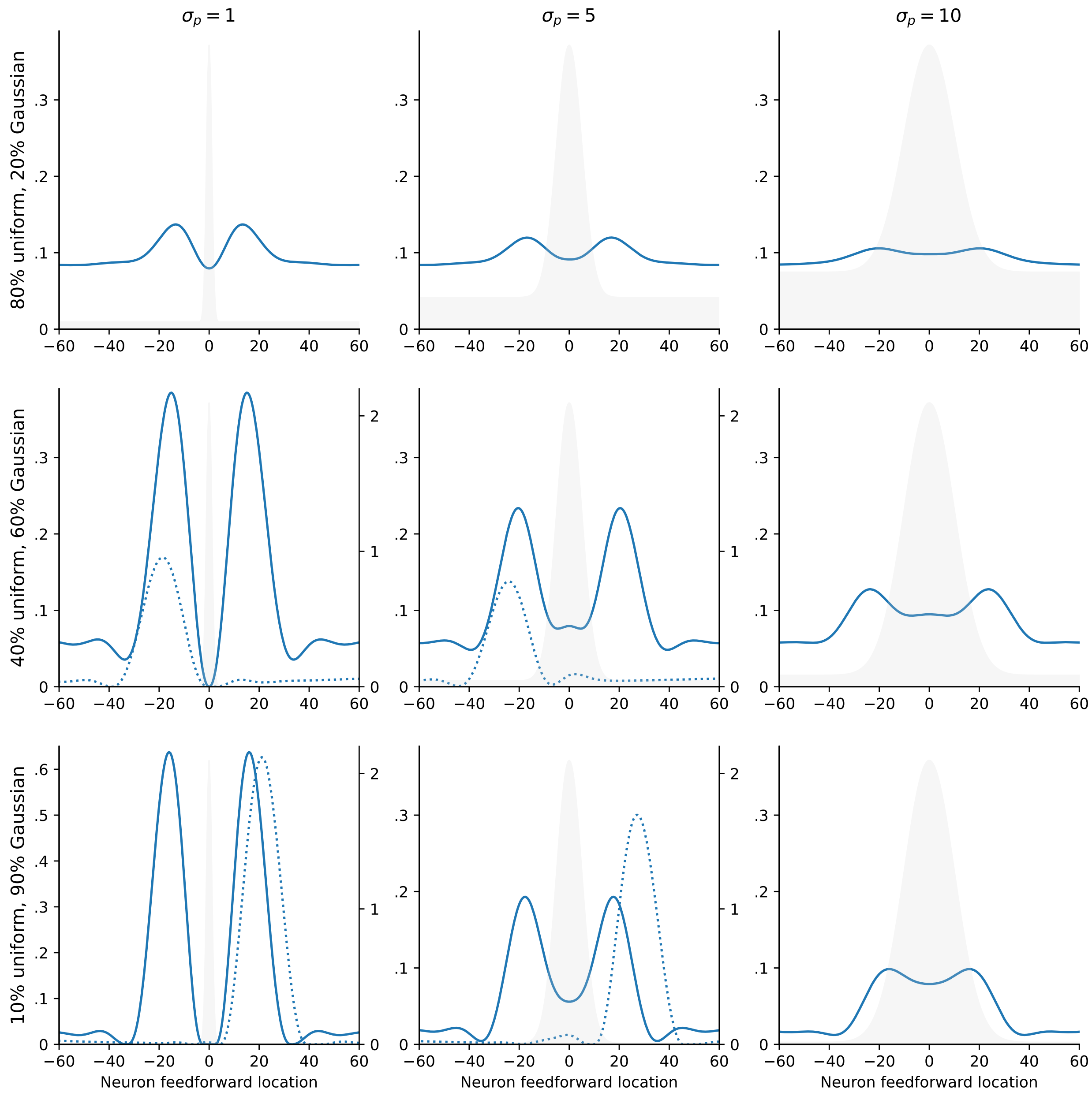


**Fig. S4: Optimal gains with various priors.** Optimal gains of the network, with mixture priors with a uniform component and a Gaussian component in the proportions 80/20 *(top row)*, 40/60 *(middle row)*, and 10/90 *(bottom row)*, and a width parameter of the Gaussian component equal to 1 *(first column)*, 5 *(second column)*, and 10 *(third column)*. The grey shaded area shows the prior. In some cases the optimal gain profile has only one global maximum, on the flank of the peak (dotted lines, right-hand-side axis). In these cases we also show the gain profile when optimized with the additional condition that it be symmetric around the peak center.

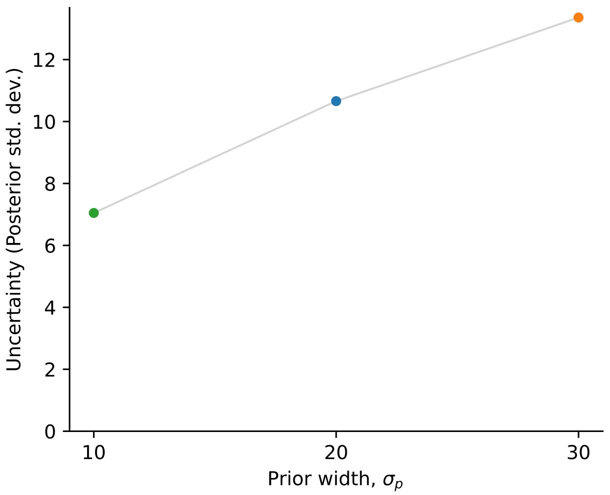


**Fig. S5: Uncertainty (posterior standard deviation) as a function of prior width.**

### Quality of approximations

We compare the approximations of various quantities that we have used in our study to their correct values. Unless otherwise noted, the various parameters have the same values as in the main text.

Figure S6 shows the eigenvalues of $W$ together with their approximations (Eq. S2), for different values of $\sigma_{rec}$. Figures S7 and S8 show the elements of the matrix $M - I$, and their approximations (Eqs. S4-S7). Figure S9 shows the covariance matrix $\Sigma$ solution of the Lyapunov equation, along with its approximations (Eqs. S24, S25). Figure S10 shows the effective locations and width as a function of the feedforward locations, the effective rates $r_i(s)$ as a function of $s$, and the square-root of the mean squared error, along with their approximations.

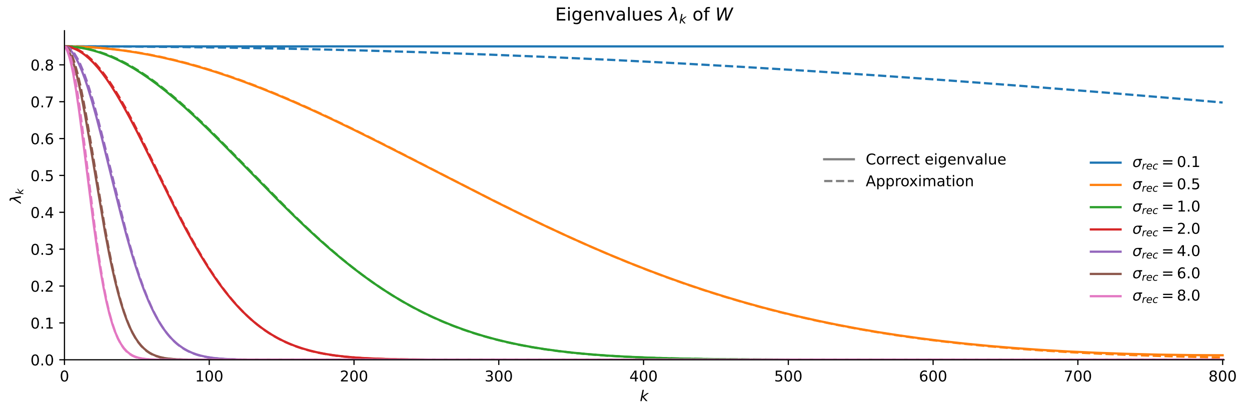


**Fig. S6: Eigenvalues $\lambda_k$ of $W$.**

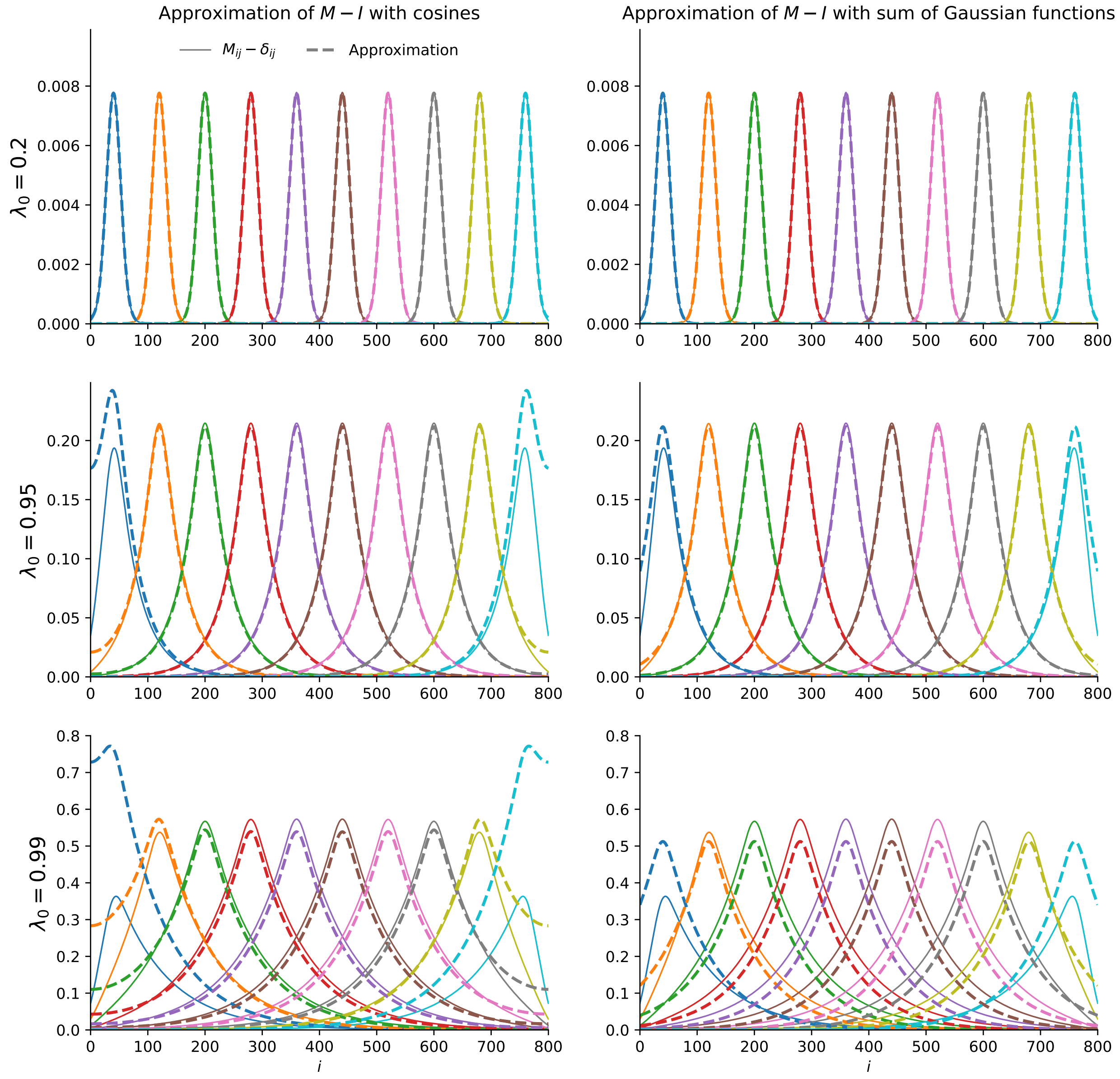


**Fig. S7: Elements of matrix $M - I$ and approximations with cosines and with sums of Gaussian functions.** Each line represents for a column $j$ the elements $M_{ij} - \delta_{ij}$, indexed by row $i$. *Left:* approximation using cosines (Eq. S4). *Right:* approximation using a sum of Gaussian functions (Eq. S5). *Top to bottom row:* $\lambda_0 = 0.2, 0.95$, and $0.99$.

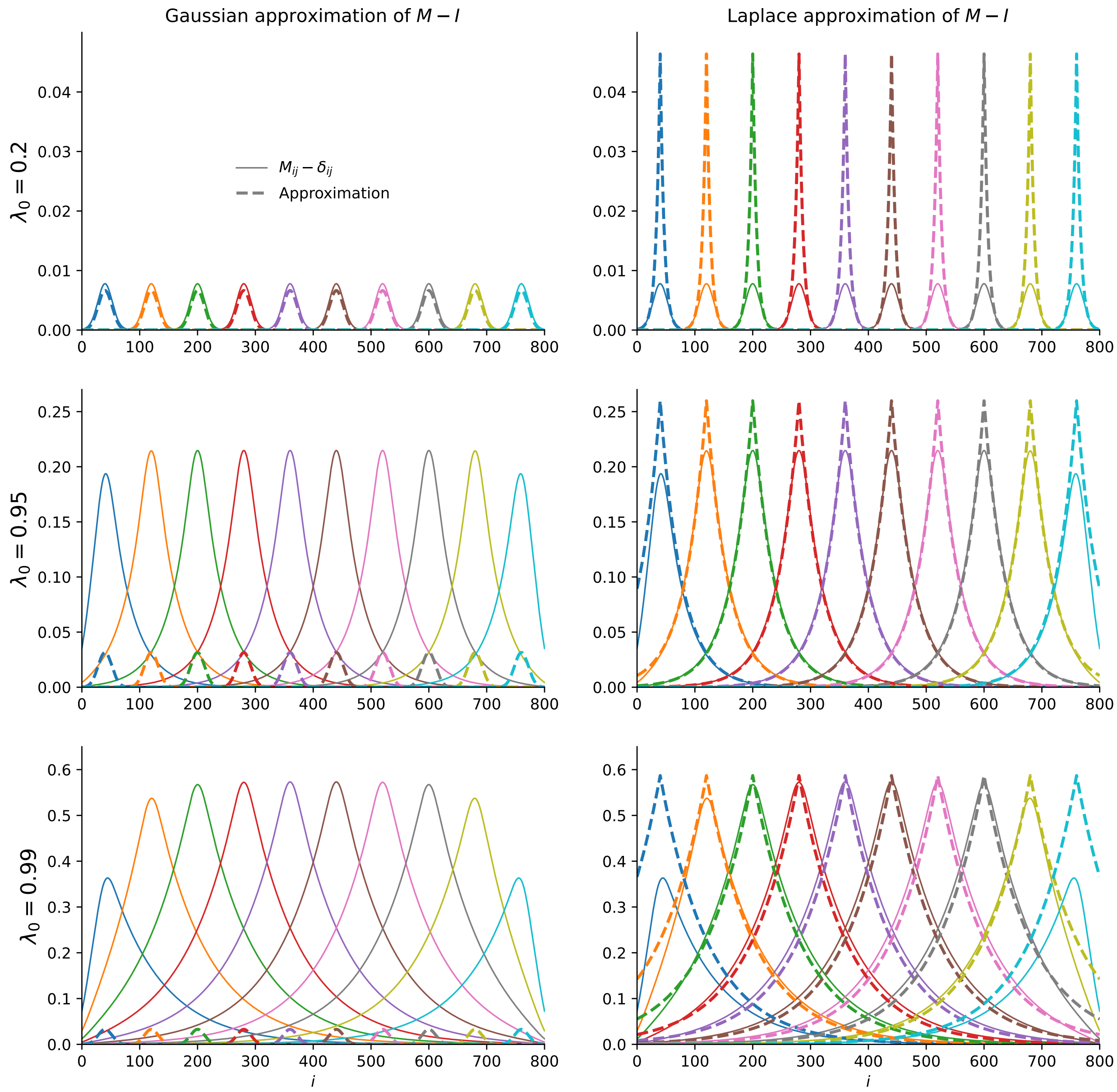


**Fig. S8: Elements of matrix $M - I$ and Gaussian and Laplace approximations.** Each line represents for a matrix column $j$ the elements $M_{ij} - \delta_{ij}$, indexed by matrix row $i$. *Top to bottom row:* $\lambda_0 = 0.2, 0.95$, and $0.99$. *Left:* Gaussian approximation (Eq. S6). *Right:* Laplace approximation (Eq. S7). For small $\lambda_0$, the Gaussian approximation is good (top left); for larger $\lambda_0$, the Laplace approximation is preferable (bottom right).

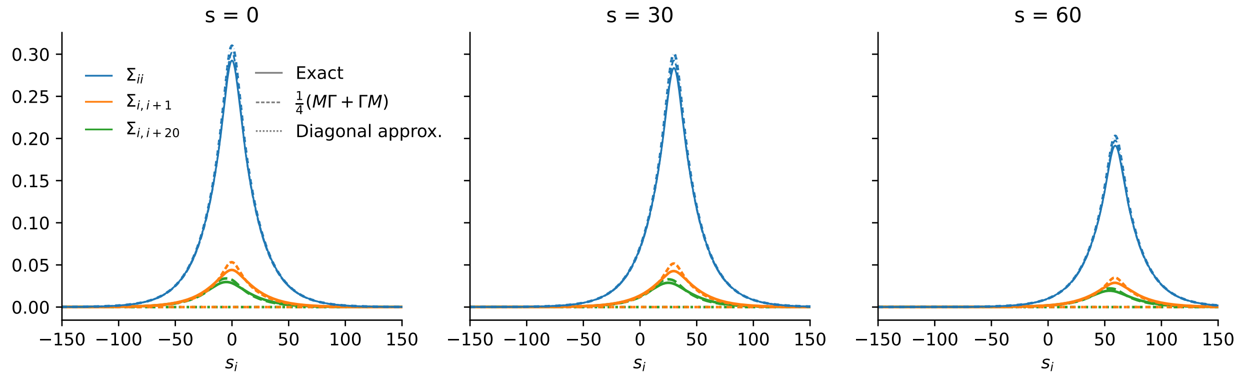


**Fig. S9: Covariance matrix $\Sigma$.** For $s = 0$ (left panel), 30 (middle panel), and 60 (right panel), with the network optimized for the Wide prior ($\sigma_p = 30$), values on the diagonal of the covariance matrix $\Sigma$ (blue lines), of the off-diagonal elements with offset 1 (orange lines), and the off-diagonal elements with offset 20 (green lines), with the covariance matrix solution of the Lyapunov equation (solid lines), the approximation $\frac{1}{4}(M\Gamma + \Gamma M)$ (dashed lines), and the diagonal approximation (dotted lines).

### Recurrent network simulations

Fig. S11 compares the empirical rates obtained by simulations, to the analytical rates (Eq. 2).

### Behavioral data

#### Effect of preceding prior

Figure S12 shows the subjects' variance as a function of the prior width, splitting trials as a function of whether the preceding prior was narrower ('Widening'), the same ('Repeated'), or wider ('Narrowing') than that in the current trial. The differences between the variances in the Widening or Narrowing trials and those in the Repeated trials are not significant.

#### Response times

##### Prior cue

At the onset of each trial the subjects were presented with the prior for this trial. This information was shown for a minimum of one second; after this delay subjects could proceed to the next step (the stimulus). The subjects' median viewing time was on average 1,650ms (sd: 699) in the Narrow condition, 1,690ms (sd: 870) in the Medium condition, and 1,673ms (sd: 720) in the Wide condition. The differences are not significant (paired t-tests p-values $> 0.25$).

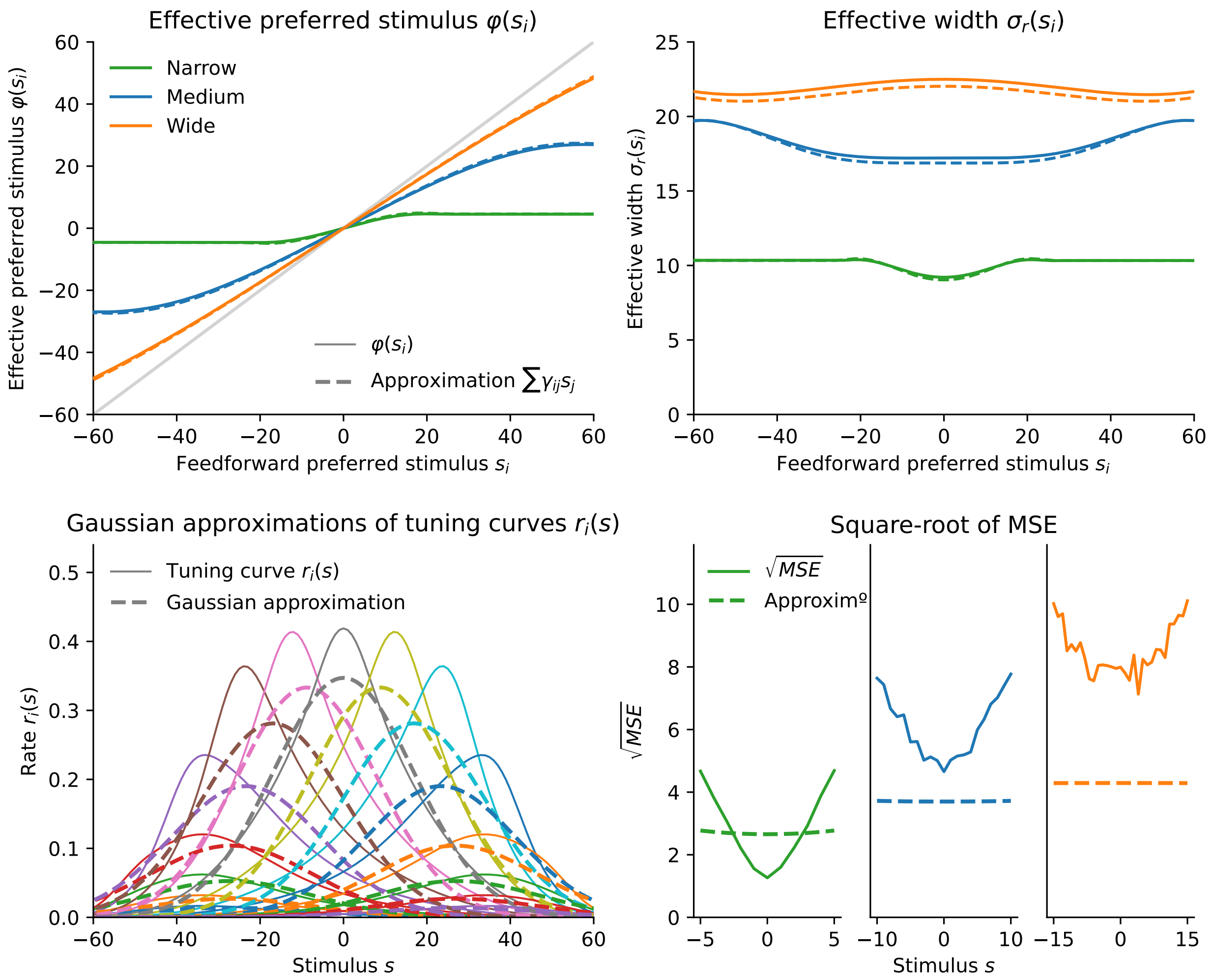


**Fig. S10: Effective tuning curves and mean squared error.** These results were obtained with optimal gains for each prior. *Top left:* Effective locations $\varphi(s_i)$ and their approximations as weighted averages (Eq. 5) as a function of the feedforward locations, $s_i$. *Top right:* Effective tuning-curve widths $\sigma_r(s_i)$ and their approximations (Eq. 6). *Bottom left:* Effective tuning curves $r_i(s)$, and their Gaussian approximations (Eq. 8), with the Medium prior. *Bottom right:* Square-root of the mean squared error (MSE) of the decoder as a function of the stimulus, $s$. The approximation corresponds to the integrand of $L$ (Eq. 14), without the prior.

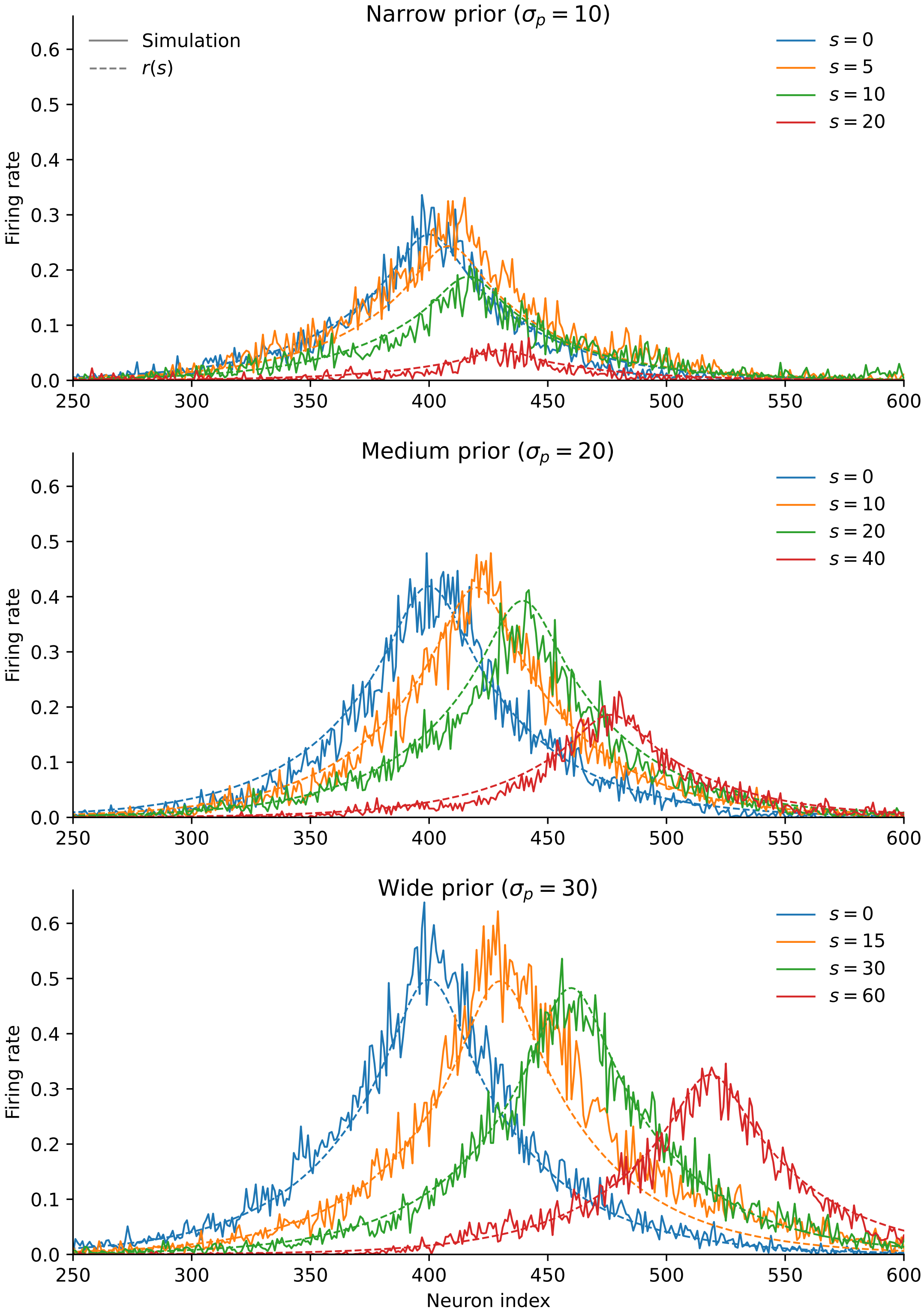


**Fig. S11: Empirical and analytical rates.** With the gains of the networks optimized to each of the three priors, empirical rates obtained with simulations of the recurrent network (solid lines), and analytical rates (Eq. 2; dashed lines), for different values of the stimulus $s$.

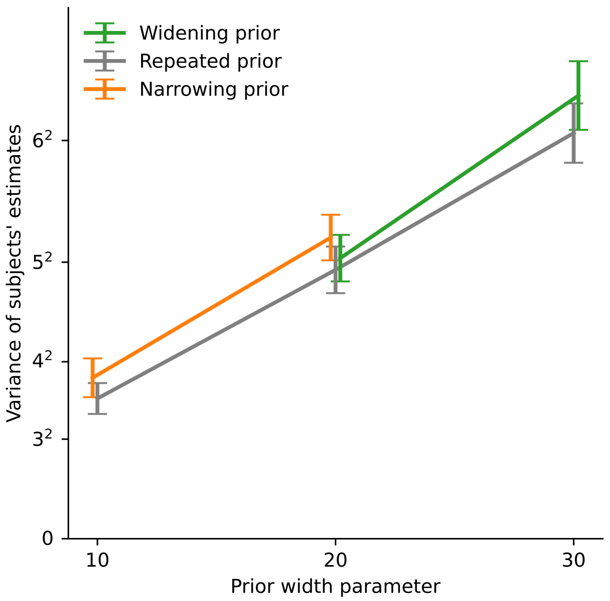


**Fig. S12: Variance vs. prior width as a function of prior history.** 'Widening prior' pools trials in which the prior is wider than that in the preceding trial (i.e., Narrow to Medium, Narrow to Wide, and Medium to Wide). 'Narrowing prior' pools trials in which the prior is narrower than that in the preceding trial (i.e., Wide to Medium, Wide to Narrow, and Medium to Narrow). 'Repeated prior' pools trials in which the prior is the same as in the preceding trial.

##### Number estimates

Subjects provided their responses using a slider. Their response times increased with the prior width: the median time was on average 3,020ms (sd: 1,101) in the Narrow condition, 3,153ms (sd: 1,054) in the Medium condition (a significant increase: paired t-test $t(58) = 2.9, p = 0.0052$), and 3,396ms (sd: 1,221) in the Wide condition (a significant increase from the Medium condition: paired t-test $t(58) = 5.5, p = 7.6 \times 10^{-7}$). We also record the slider 'trajectory', i.e., all the selected value until the final, submitted one. The mean trajectory length was 2.9 (sd: 1.8) in the Narrow condition, 3.5 (sd: 2.5) in the Medium condition (paired t-test $t(58) = 3.7, p = 0.0004$), and 4.8 (sd: 3.8) in the Wide condition (compared with Medium, paired t-test $t(58) = 6.7, p = 1.1 \times 10^{-8}$). Thus presumably subjects take longer with the wider priors because they go through more values on the slider before finding the desired number.

#### Model of Gaussian encoding

To investigate the sensitivity of subjects' responses to the prior width, and to examine the case of a Bayesian observer whose encoding does not change across conditions, we fit the following model of Gaussian encoding [10]. In this model, the presentation of a number $x$ elicits a noisy signal $r$ drawn from a Gaussian distribution centered on $x$ and with variance

$\nu^2 w^{2a}$, i.e.,

$$r|x \sim N(x, \nu^2 w^{2a}), \tag{S79}$$

where $w$ is the width of the prior, $\nu^2$ is the 'baseline' variance of the noise, and the exponent $a \geq 0$ is a parameter that controls the dependence of the imprecision on the prior width. Upon receiving the signal $r$ one can derive the Bayesian posterior mean, $x^*(r) = \mathbb{E}[x|r]$, which would in our task maximize the expected reward. Our model includes motor noise: we assume that the response $\hat{x}$ is drawn from a Gaussian distribution centered on $x^*(r)$, with variance $\sigma^2 > 0$, and truncated to the slider interval.

The case $\nu = 0$ corresponds to a 'motor-noise-only' model, that features no internal cognitive noise. If $\nu > 0$ and $a = 0$, then there is cognitive noise in addition to motor noise, but the scale of this cognitive noise does not change with the prior width. If $a > 0$, wider priors result in larger imprecision. In particular, Reference [10] presents a model of endogenous precision which predicts that for an estimation task, $a = 1/2$, and the study provides empirical support for this value of the exponent $a$. This model predicts that the variance increases as a linear function of the prior width. An alternative model would be the specification $a = 1$. This 'normalization' model predicts that the variance increases as a linear function of the square of the prior width.

We fit these models with different values of $a$, letting $\nu$ and $\sigma$ be free parameters for each subject (except when we enforce $\nu = 0$). The model with $a = 1/2$ best fit our data. Specifically, it yields the lowest Bayesian Information Criterion (BIC [71]; see Table 1). The motor-noise-only model results in the highest BIC (i.e., poorest fit), and the BIC of the model with $a = 1$ is higher by 420 than that of the model with $a = 1/2$, indicating very strong evidence in favor of the hypothesis $a = 1/2$. The BIC of the model with $a = 0$ is higher by 3.3 (than with $a = 1/2$), indicating moderate evidence in favor of the model with $a = 1/2$. We thus fit another model in which $a$ is a free parameter allowed to be different for each subject. This model yields a BIC higher by 129 than that of our best-fitting model, suggesting that the model with $a = 1/2$ provides a better and more parsimonious account of the data than assuming that the exponent $a$ is different for all the subjects. We can however look at the best-fitting values of $a$, across subjects. We find that for 27% of subjects, the best-fitting exponent is $a = 0$, while for the remaining majority of subjects, the best-fitting $a$ is on average 0.45 (sd: 0.29), a value not far from 1/2. We conclude that although some subjects may maintain the same encoding across different priors, a large fraction of subjects seem to adapt their encoding imprecision to the prior width.

| Model | BIC |
|---|---|
| $\nu = 0$ | 61170.35 |
| $a = 0$ | 58486.46 |
| $a = 1/2$ | *58483.16 |
| $a = 1$ | 58903.57 |
| Free $a$ | 58613.06 |

**Table 1: Gaussian models BICs.** The asterisk indicates the lowest BIC.
